# The contribution of PARP1, PARP2 and poly(ADP-ribosyl)ation to base excision repair in the nucleosomal context

**DOI:** 10.1101/2020.10.05.325944

**Authors:** M.M. Kutuzov, E.A. Belousova, T.A. Kurgina, A.A. Ukraintsev, I.A. Vasil’eva, S.N. Khodyreva, O.I. Lavrik

**Affiliations:** Institute of Chemical Biology and Fundamental Medicine, SB RAS, Novosibirsk, Russia; Novosibirsk State University, Novosibirsk, Russia

**Keywords:** PARP1, PARP2, nucleosome, base excision repair, poly(ADP-ribosyl)ation, APE1, DNA polymerase β, DNA ligase IIIα, XRCC1

## Abstract

The repair processes regulation including base excision repair (BER) is implemented by a cellular signal PARylation catalysed by PARP1 and PARP2. Despite intensive studies, it is far from clear how BER is regulated by PARPs and how the roles are distributed between the PARPs. Here, we investigated the effects of PARP1, PARP2 and PARylation on activities of the main BER enzymes (APE1, Polβ and LigIIIα) in combination with XRCC1 in the nucleosomal context. We constructed nucleosomes with midward- or outward-oriented damage. It was concluded that in most cases, the presence of PARP1 leads to the suppression of the activities of APE1, Polβ, and to a lesser extent LigIIIα. PARylation by PARP1 attenuated this effect to various degrees. PARP2 had an influence predominantly on the last stage of BER: DNA sealing. Nonetheless, PARylation by PARP2 led to Polβ inhibition and to significant stimulation of LigIIIα activities in a NAD^+^-dependent manner.

## INTRODUCTION

Genome stability in higher eukaryotes is maintained by the activity of several DNA repair pathways selective to the type of DNA damage (1). DNA lesions that do not induce strong distortions in the double helix of DNA structure are eliminated by base excision repair (BER) (2, 3). Even though this process was discovered a long time ago, its regulation is still actively studied.

BER includes the following main stages: DNA damage recognition, excision of the damaged base, incision of the sugar-phosphate backbone, gap processing (including the incorporation of dNMP) and DNA ligation. At the initiation stage, a lesion-specific DNA glycosylase, which could be a mono- or bifunctional enzyme, recognises a damaged heterocycle and hydrolyses the N-glycosidic bond thereby forming an apurinic/apyrimidinic (AP) site in the DNA duplex. In addition, the emergence of an AP site in DNA could be a consequence of spontaneous hydrolysis of the N-glycosidic bond (2). Bifunctional DNA glycosylases cleave DNA with the formation of 3′-phospho-α,β-unsaturated aldehyde and 5′-phosphate ends or 3′- and 5′-phosphate ends depending on the reaction mechanism (β- or β/δ-elimination) (4).

AP sites are predominantly cleaved via hydrolytic mechanisms resulting in a single-strand break containing 5′-deoxyribose phosphate and 3′-hydroxyl groups. In mammalian cells, this reaction is mostly catalysed by AP endonuclease 1 (APE1) (5). Moreover, APE1 cleaves 3′-phospho-α,β-unsaturated aldehyde thereby producing a 3′-hydroxyl group. Thus, APE1 is the main enzyme that creates single-strand breaks with 3′-hydroxyl groups for the synthetic stage of the BER process.

The canonical short-patch pathway of BER proceeds in the following way. Continuing our review of BER, at the next stage, the single-nucleotide gap is filled by sequential removal of 5′-deoxyribose phosphate by 5′-deoxyribose phosphate (dRP) lyase activity and incorporation of dNMP via the nucleotidyl transferase activity of DNA polymerase β (Polβ). Finally, the single-strand break is sealed by the activity of DNA ligase IIIα (LigIIIα) in complex with XRCC1.

Accurate BER functioning strongly depends on the regulation of each stage of the process. There are several proteins responsible for the BER regulation: XRCC1, PARP1 and PARP2 (6–9). Moreover, the regulation of different repair processes during the DNA damage response is driven by the emergence of special cellular signals; one of them is poly(ADP-ribosyl)ation (PARylation) (10). The metabolism of poly(ADP-ribose) (PAR) in the nucleus is carried out by nuclear poly(ADP-ribose) polymerases (PARPs) 1 and 2, poly(ADP-ribose)glycohydrolase (PARG) and other PAR erasers and regulates various nuclear processes such as DNA repair, DNA replication, gene expression, chromatin dynamics, cell death and mitotic progression (11). Using NAD^+^ as a substrate, several members of the PARP family can modify themselves (by PAR) and protein partners to modulate their functioning. PARP1 is the most abundant and active PARP, is strongly activated in response to DNA damage and synthesises most of PAR after genotoxic or oxidative stress (12–14). Additionally, recent studies clarified the functions of PARP1 in normal proliferating cells during the S phase: Okazaki fragment maturation and regulation of replication fork speed (15, 16). Hypersensitivity to small-molecule PARP inhibitors is reported for the cells lacking efficient homologous-recombination–mediated repair (17, 18). Mice devoid of PARP1 or PARP2 are viable but exhibit sensitivity to genotoxic agents, thus pointing to the involvement of both proteins in DNA repair (19, 20). Nonetheless, double deletion of PARP1 and PARP2 causes early embryonic lethality in mice, indicating that each of these two proteins can partly compensate for the absence of the other one (21).

The interaction of PARP1 with various BER DNA intermediates as well as the influence of PARP1 on the activity of key BER proteins have been analysed by diverse methods (6, 22-28). These data show an interaction of PARP1 with an earlier intermediate of BER: the AP site. Additionally, it was documented the interaction of PARP1 with the central BER intermediate formed by the AP site– cleaving activity of APE1 as well as the influence of PARylation on the activity of Polβ and the other BER enzymes in short-patch and long-patch BER pathways (22-26).

Hanzlikova and co-authors have supposed overlapping roles of PARP1 and PARP2 in the implementation of BER and/or single-strand break repair processes (29). They have found that the activity of PARPs is important for recruiting the key repair proteins XRCC1 and PNKP to chromatin lesions.

In addition, there are some data on the affinity for PARP2 and its activation by various DNA substrates; thus, it is clear that this enzyme can be involved in later stages of BER (8, 14, 30-32). Moreover, the interactions of PARP2 with BER proteins XRCC1, Polβ and LigIIIα have been demonstrated in HeLa cells (33). Recently, the specificity of the interaction of PARP1 and PARP2 with various BER intermediates was analysed by atomic force microscopy (AFM) using long DNA duplex substrates, and the data speak in favour of PARP2 contribution to the later stages of BER (31). Nevertheless, the final function of PARP2 in this process is not clear.

To date, the main biochemical steps of BER have been investigated fairly well. The bulk of experimental data on the activity of BER proteins has been obtained by means of naked DNA. On the other hand, DNA in eukaryotic cells is condensed to different degrees depending on a cell cycle phase and DNA region (34, 35). Therefore, most of attention is now focused on the regulation of DNA repair in the chromatin context. There are many studies revealing the efficiency of individual BER stages by means of nucleosomes, but there is no information about the roles of nuclear PARPs and their activities in this process (reviewed in 36, 37). Accordingly, it is interesting how BER is regulated by the nuclear PARPs and how the roles are distributed between different PARPs. Therefore, the aim of this study was to investigate the effects of PARP1, PARP2 and PARylation on principal BER stages using key BER proteins: APE1, Polβ, XRCC1 and LigIIIα at the nucleosomal level (Fig. 1C). On the basis of literature data concerning the involvement of PARP1 and PARP2 in BER and our present results relating to the functional interactions of these proteins in BER on nucleosome core particles (NCPs), we discuss a hypothetical model of the contributions of PARP1 and PARP2 to BER. We propose that different stages of the BER process are regulated by PARP1 and PARP2, and the effect depends on PARylation.

**Figure 1.**
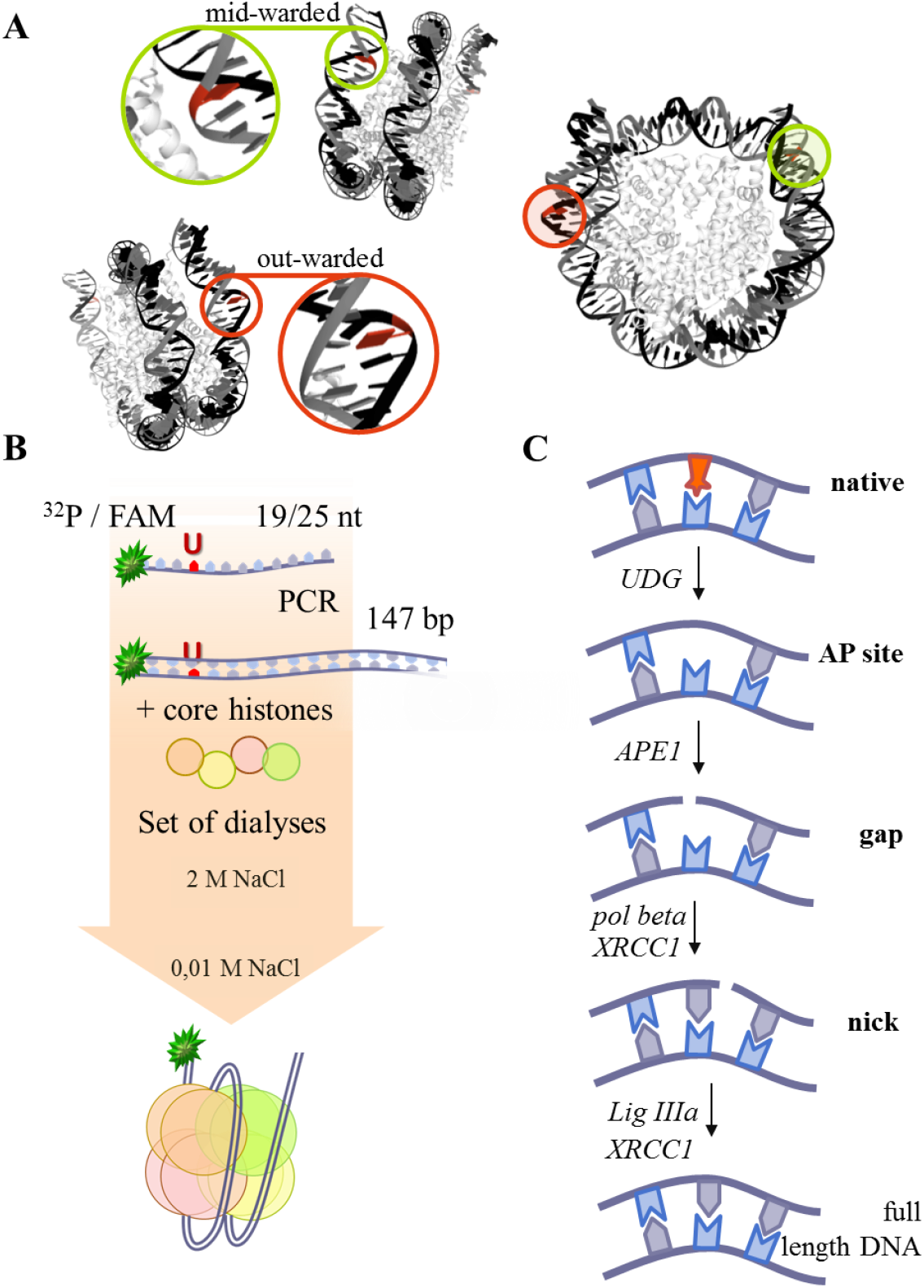
(A) Nucleosome structure pictured according to (43). The red nucleobases depict the positions of midward- or outward-oriented lesions. (B) The scheme of preparation of different types of NCPs. The coloured line corresponds to the DNA strand, whereas the green asterisk to the 5′[^32^P] or FAM label. (C) Stages of BER with the participating enzymes and substrates tested in this study.

## RESULTS

### PARP1 or PARP2 efficacy in the interaction with and the activation by damaged NCP

The influence of PARPs and/or of PARylation on the BER process in the nucleosomal context may be complicated due to the complex nature of the substrate. The lowest level of DNA compaction in chromatin is the NCP, which consists of a 147 bp DNA wrapped around the histone octamer core: two of H2A, H2B, H3 and H4 each; this compaction level affects DNA–protein interactions (38). It should be noted that with this DNA wrapping, the accessibility of heterocyclic bases to other DNA–protein interactions varies greatly with the DNA sequence and nucleotide position (39-41). There is a special nomenclature for the spatial orientation of the bases: ‘inward’ if the base is turned towards the histone core, and ‘outward’ if the base is facing out (36, 42). Some bases may have an intermediate, ‘middle’, orientation (40). The orientation of the base directly affects the nature of the DNA–protein interactions and therefore functional activity of the enzymes. Outward-oriented nucleobases are the most accessible to repair enzymes; on the contrary, inward-oriented nucleobases are the least accessible for interactions. In an assay of DNA–protein interaction, the use of substrates with an inward orientation of damage will complicate the assessment of the potential contribution of the positional effect, owing to steric hindrance. For these reasons, to assess the effects of PARP1, PARP2 and PARylation on the activity of BER enzymes, NCPs with outward- or midward-oriented damage were tested in our study (Fig. 1A).

To determine the contribution of the interaction of PARP1 or PARP2 with DNA or histone components of an NCP in the BER process, the affinity of the PARPs for native, AP-NCP or gap-NCP was evaluated by fluorescence measurements (Table 1, Fig. S2, A–E).

**Table 1.**
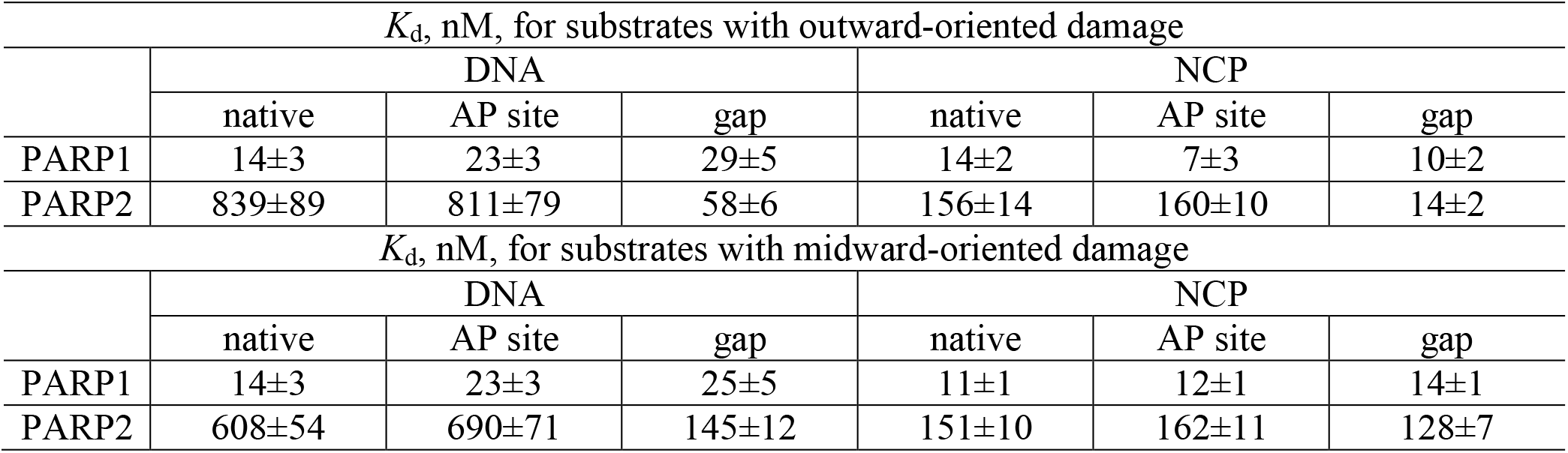
The affinity of PARP1 and PARP2 for native, AP site–containing or gap-containing substrates as determined by fluorescence anisotropy measurements

| $K_d$, nM, for substrates with outward-oriented damage | | | | | | |
| --- | --- | --- | --- | --- | --- | --- |
|  | DNA |  |  | NCP |  |  |
|  | native | AP site | gap | native | AP site | gap |
| PARP1 | 14±3 | 23±3 | 29±5 | 14±2 | 7±3 | 10±2 |
| PARP2 | 839±89 | 811±79 | 58±6 | 156±14 | 160±10 | 14±2 |
| $K_d$, nM, for substrates with midward-oriented damage | | | | | | |
|  | DNA |  |  | NCP |  |  |
|  | native | AP site | gap | native | AP site | gap |
| PARP1 | 14±3 | 23±3 | 25±5 | 11±1 | 12±1 | 14±1 |
| PARP2 | 608±54 | 690±71 | 145±12 | 151±10 | 162±11 | 128±7 |

PARP1 generally proved to have high affinity for the NCP as well as for DNA. It is noteworthy the lacking of strong differences in *K*_d_ between the complexes of PARP1 with the NCP or DNA that could be explained by high affinity of the protein for DNA blunt ends. In our case, the interaction of PARP1 with a specific DNA damage site such as an AP site or gap likely can be masked by an interplay of this protein with nearby blunt end of the DNA duplex.

The character of the PARP2 interaction with DNA or NCP is more complicated than that of PARP1. Our data uncovered low affinity of PARP2 for an undamaged or AP site–containing naked DNA; for comparison, the affinity of PARP2 for gapped DNA was higher by one order of magnitude. Additionally, PARP2 manifested dramatically higher affinity for an NCP than for naked DNA. PARP2 was found to have one order of magnitude higher affinity for outward-oriented damage in an NCP as compared to a midward-oriented one. Moreover, PARP2 prefers the gap-containing DNA duplex in the nucleosomal context, as is the case for naked DNA (Table 1, Fig. S2, A–E).

The interaction of PARP1 or PARP2 with different types of outward-oriented NCP was additionally tested by the EMSA (Table S1, Fig. S2H and S2I). The results illustrate basic patterns of interactions for both PARPs as determined by the fluorescence measurements. Higher affinity of PARP1 for an NCP was demonstrated. Moreover, the *K*_d_ values obtained by the EMSA revealed the affinity of PARP1 for the gapped substrate. The patterns identified during the interaction of PARP2 with different NCPs by the fluorescence measurements were confirmed by the EMSA.

It is known that the PARP activation efficiency is poorly correlates with the affinity of the protein for the substrate, but is largely determined by the cooperative interaction of the molecule domains upon interaction with the substrate (44, 45). In this work, PARP1/PARP2 activation efficacy was studied in the presence of native or containing various damage types either DNA or NCP using [32P]radioactive NAD^+^ incorporation into growing ADP-ribose polymer chain. The data are presented in Fig. 2.

**Figure 2.**
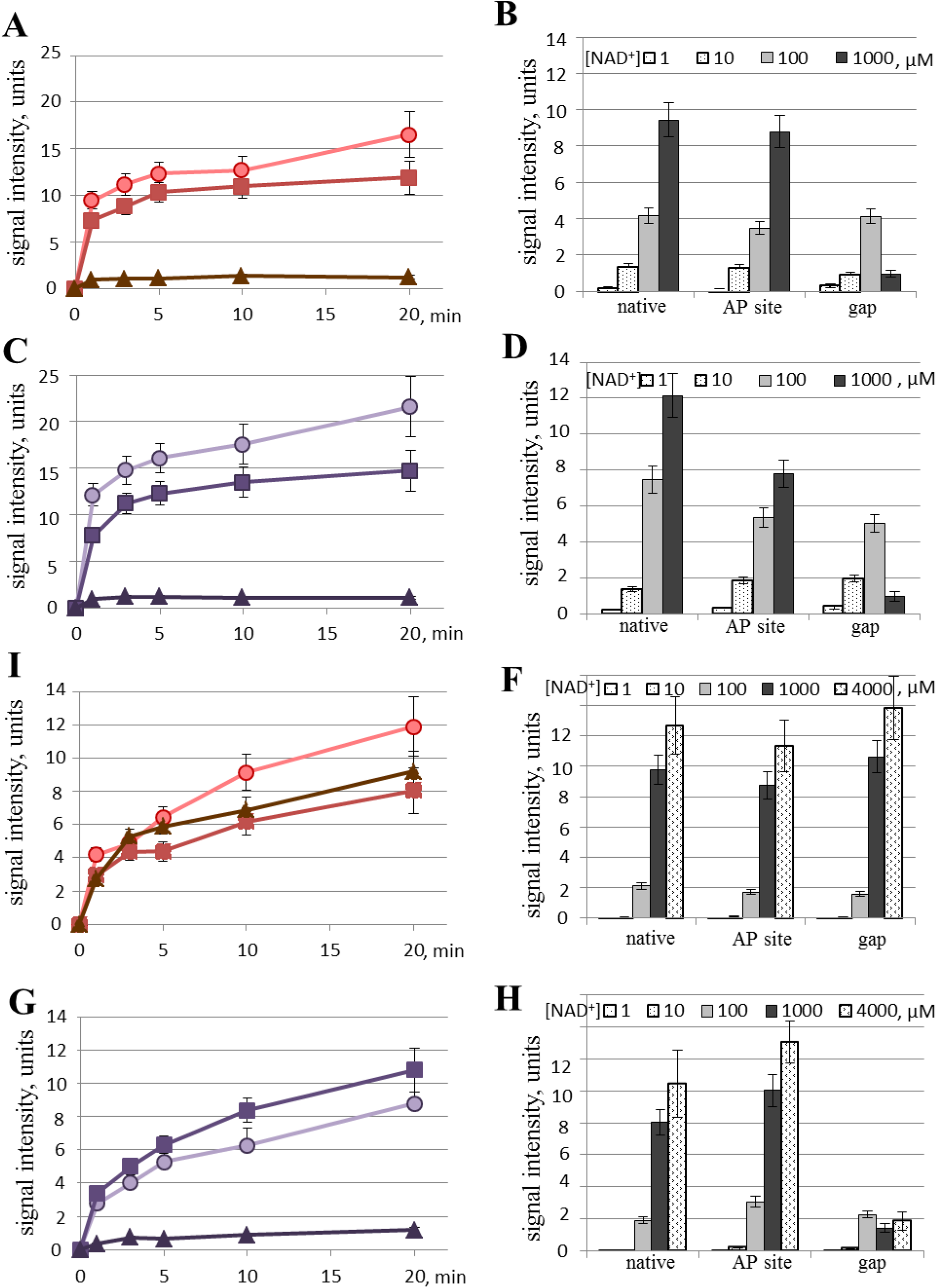
The activation of PARP1 (A-D) and PARP2 (I-H) by native (circles), AP (squares) site and gap-containing NCP (triangles) (C, D, G, H) or DNA (A, B, I, F) as measured by as the amount of an poly(ADP-ribose) synthesised by PARP1/PARP2 using [^32^P]labelled NAD^+^ as a precursor. The graphs in panels (A), (C), (I) and (G) represent the accumulation kinetics of PAR at the 1000 μM NAD^+^, the histograms in panels (B), (D), (F), (H) represent the total amount of PAR synthesised by PARP1 for 1 min and by PARP2 for 3 min at NAD^+^ concentrations 1-1000 μM or 1-4000 μM, respectively. In all the graphs, the experimental data on NCP substrates correspond to the red or yellowish-green curves; the DNA substrates are characterised by black, reddish-violet or violet curves. The data are presented as an average of at least three independent experiments and showed the mean values ± SD. The reactions were carried out as described in ‘Materials and Methods’.

In general, the data obtained using mid- and outwarded NCP or subsequent DNA were in a good correlation with each other in the case of both PARPs. Upon PARP1 activation by the NCP the efficiency of radioactive PAR synthesis was higher than by naked DNA, especially in the absence of the internal lesions (Fig. 2A and 2C). At the same time, internal lesion in the DNA structure caused a decreasing of the total amount of synthesized PAR as NAD^+^ concentration was increased up to 1000 μM, regardless of the presence of a nucleosomal structure; the maximal effect of PAR synthesis inhibition was obtained upon PARP1 interaction with gapped substrates (Fig. 2B and 2D).

The efficiency of PARP2 activation upon the interaction with nucleosome was approximately the same with the activation by free DNA; the only exception was gap-NCP (Fig. 2I and 2G). The total amount of PAR synthesised in the presence of a gap in the nucleosome structure was decreased significantly too (Fig. 2F and 2H). This effect seems especially remarkable in the nucleosomal context because the amount of PAR synthesised in the presence of naked DNA did not change upon the appearance of the lesions.

In our study intended to examine the contribution of PARylation to the binding of a nucleosome by PARP1 or PARP2, real-time fluorescence anisotropy measurements of PARPs’ activity in the presence of NAD^+^ were performed (46). These experiments were carried out with NCP substrates containing an outward-oriented lesion (Table 2, Fig. S2F and S2G). Here, *K*_M_ values are presented as a complex constant reflecting the processes of PARP binding to the NCP, NAD^+^ binding by PARP’s NAD^+^- binding site, subsequent conformational changes of the PARP–NCP complex, catalysis of PARylation, the dissociation of the PARylated PARP–NCP complex, and size changes of the PARP–NCP complex driven by PAR synthesis. The obtained *k*_obs_ values likely reflect an overlay of the following effects: nucleosome refolding and dissociation of the PARylated PARP-NCP complex.

**Table 2.**
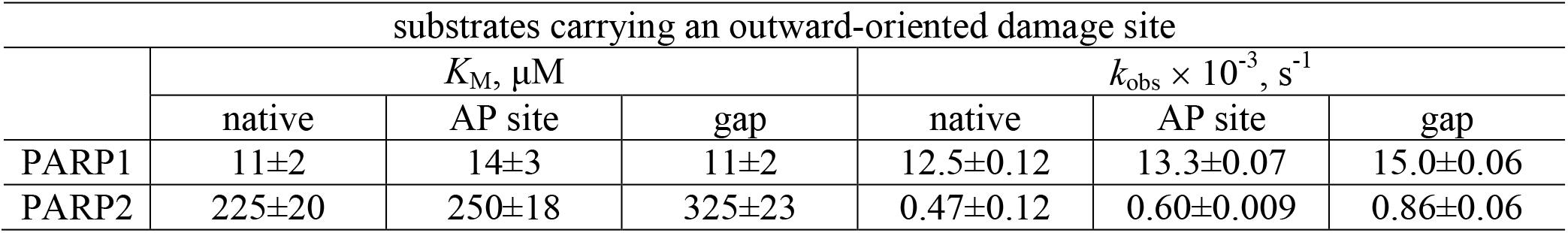
K_M_ and k_obs_ values for PARylation by PARP1 and PARP2 during the interaction with a native NCP, AP-NCP or gap-NCP according to fluorescence anisotropy measurements

Readers can see that the *k*_obs_ values under PARP2 PARylation were lower as compared to PARP1. PARP1 relatively quick dissociated as compare to PARP2. This fact may be explained by the lowered activity of PARylation catalysed by PARP2 and consequently the rate of dissociation of PARylated PARP2 from the substrate (8, 47).

Only insignificant dissimilarities in the *K*_M_ values among different NCP structures were observed for PARP1. Nevertheless, the *k*_obs_ values were growing under PARP1 PARylation in the following order: native NCP ≤ AP-NCP < gap-NCP. Taking in mind that the Kd values are relatively close, this finding can be attributed to the effective interaction of PARP1 with the blunt ends of DNA duplexes and relatively whipping dissociation of the autoPARylated protein from the substrate.

As for PARP2, all the analysed parameters are more than an order of magnitude higher as compared to PARP1. *K*_M_ for the gapped substrate is higher than that for the native or AP site–containing structure. By contrast, *k*_obs_ values increased in the following order: native NCP < AP-NCP < gap-NCP.

Therefore, our data are suggestive of the expected contribution of PARP1 at the different stages of BER. As for PARP2, its contribution is expected at later stages of the BER process, starting from the formation of gapped substrates. The character of the interaction of PARP2 with the damaged NCP reveals an impact of the interaction with nucleosomal proteins or a significant modification of the interaction with DNA in the nucleosomal context.

### The activity of APE1 and effects of PARP1 and PARP2

APE1 is one of the earlier enzymes in the general BER scheme and is the first enzyme initiating AP site repair. Numerous studies on APE1 involving both naked DNA and an NCP indicate that the nucleosomal organisation of DNA poses an obstacle for the access to the AP site. Therefore, at the beginning of our study, specific conditions were found that ensured equivalent APE1 efficiency between nucleosomes with an midward- or outward-oriented AP site (mid-NCP and out-NCP, respectively; Fig. 3). As expected, the processing of midward-oriented damage required more than 10-fold higher concentration of the enzyme. It should be noted that the stability of the nucleosomal DNA duplex proved to be extremely high even under denaturing conditions, thus giving rise to two bands of DNA on the electropherograms: corresponding to single-stranded and double-stranded 5′-FAM– labelled DNA.

**Figure 3.**
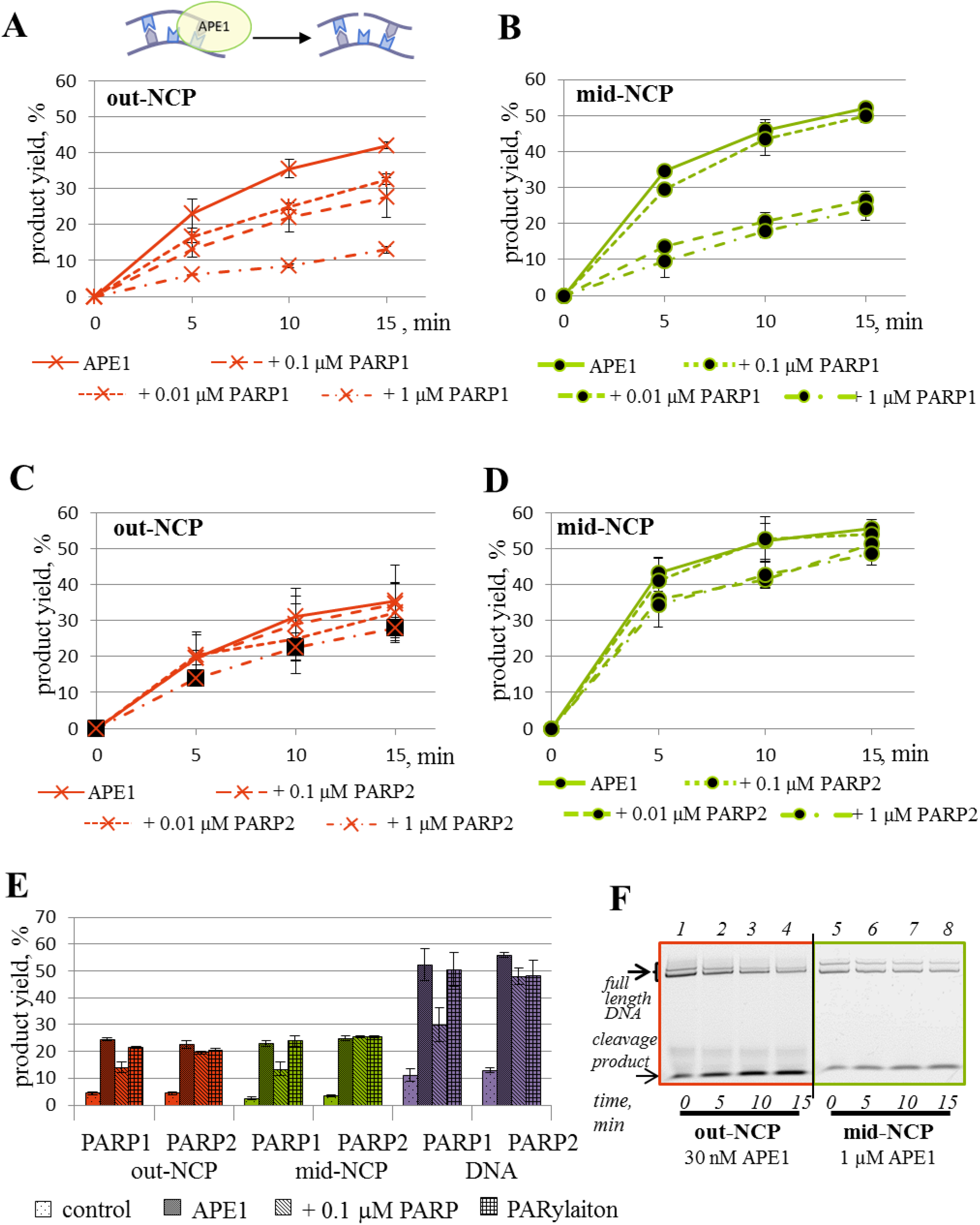
The kinetic assay of APE1 activity towards an outward- (A, C) or midward-oriented (B, D) AP site in the NCP context in the presence of PARP1 (panel A: using an NCP with outward-oriented damage, i.e. out-NCP; panel B: using an NCP with midward-oriented damage, i.e. mid-NCP) or PARP2 (panel C: using out-NCP, D: using mid-NCP). The data are presented as an average of at least three independent experiments and showed the mean values ± SD. The reactions were carried out as described in ‘Materials and Methods’. All graphs are normalised to spontaneous cleavage of an AP site under the experimental conditions. (E) The influence of PARylation catalysed by PARP1 or PARP2 on the activity of APE1 towards AP site–containing DNA or NCP-based substrates. The ‘control’ label corresponds to the spontaneous cleavage of the AP site; ‘APE1’ indicates the specific cleavage by APE1 under the reaction conditions; ‘+ 0.1 μM PARP’ denotes the specific cleavage in the presence of the indicated protein; and ‘PARylation’ represents the specific cleavage in the presence of the indicated protein and 400 μM NAD^+^. (F) Separation of the products (on a 10% denaturing polyacrylamide gel) of the reaction of 0.1 μM 5′-FAM–labelled AP-NCP with APE1 under the indicated experimental conditions. In all cases, the APE concentrations of 0.03 and 1 μM were chosen for the reaction conditions involving substrates out-NCP and mid-NCP, respectively.

At the next stage, the influence of the PARPs on the AP site cleavage was investigated. To this goal for correct detection the inhibiting or stimulating effects of PARPs and PARylation, the reaction conditions that yielded 40–50% substrate cleavage were chosen. It turned out that regardless of the damage orientation, PARP1 suppressed the activity of APE1 (Fig. 3A and 3B). The synthesis of PAR by PARP1 in the presence of NAD^+^, accompanied by the autoPARylation of PARP1, substantially restored the magnitude of cleavage of both NCP substrates by APE1 (Fig. 3E).

As for PARP2, it did not exert a significant effect on the AP site cleavage either by itself or under PARylation conditions (Fig. 3C, 3D and 3E). It is noteworthy that the PARP2 impact did depend on neither substrate organization nor rotational position of the damage.

It should be pointed out that the patterns of the influence of PARP1 or PARP2 and PARylation on the APE1 activity towards naked DNA were similar with the patterns towards out-NCP and mid-NCP, although the effects of PARP1 were stronger (Fig. 3E).

### DNA synthesis catalysed by Polβ in the presence of PARP1 and PARP2

According to the general BER mechanism, the next stage after AP site cleavage is the DNA synthesis catalysed by Polβ. In this stage, it was necessary to assess the ability of Polβ to process gap-NCP (Figs. 4A, 4B and S3). It turned out that the efficacy of DNA synthesis differed 20-fold between the substrates with an outward- or middle-ward oriented lesion [*k*_cat_ (out-NCP) ≈ 178 ± 23 × 10^−3^ s^−1^ and *k*_cat_ (mid-NCP) ≈ 9.7 ± 0.8 × 10^−3^ s^−1^]; however, *K*_M_ values were very similar [*K*_M_ (out-NCP) ≈ 1.05 ± 0.17 μM and *K*_M_ (mid-NCP) ≈ 0.79 ± 0.10 μM].

**Figure 4.**
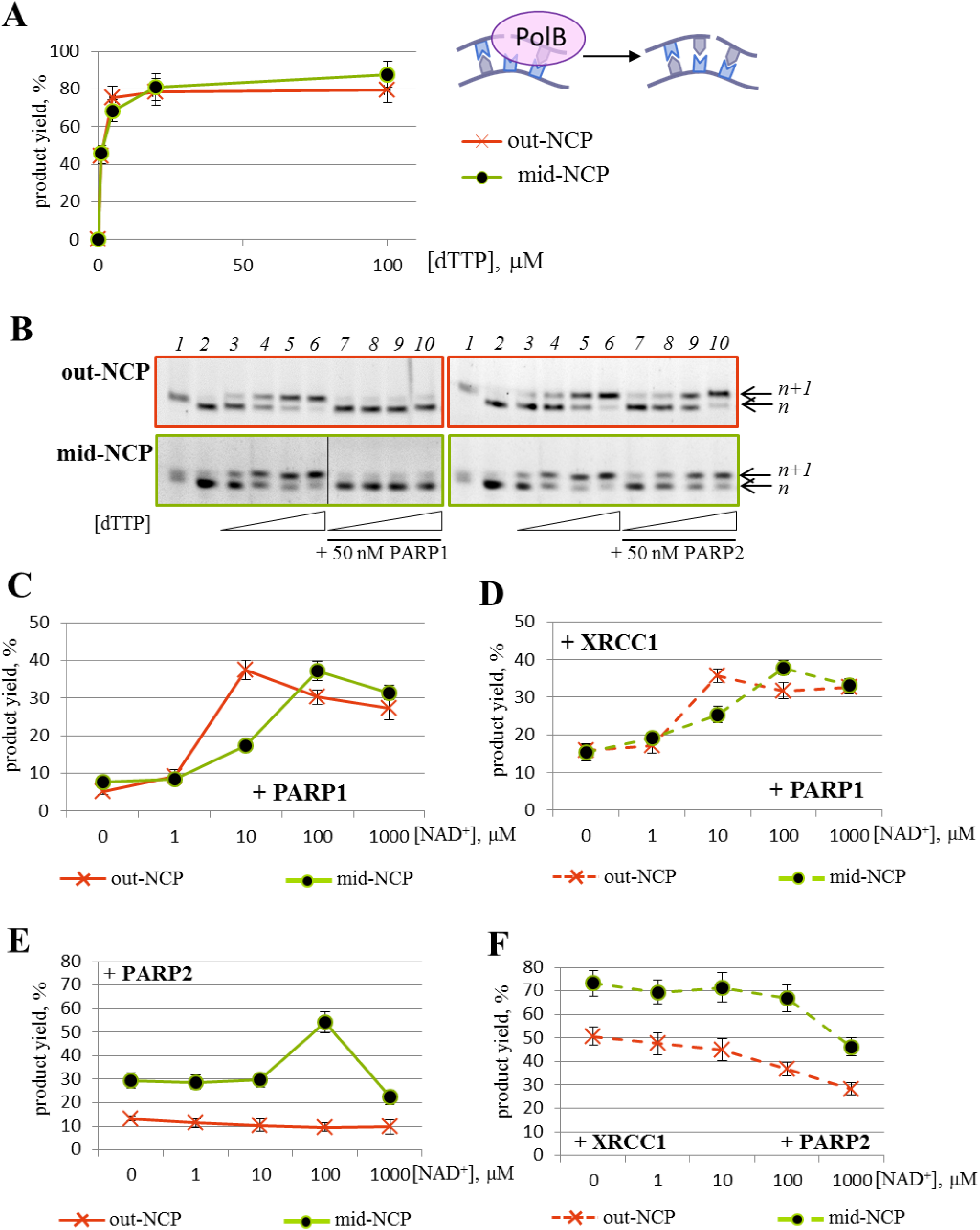
The activity of Polβ towards gap-NCP substrates by itself (A, F) and during PARylation catalysed by PARP1 (B, C) or PARP2 (D, E) in the absence (A, B, D) or presence of XRCC1 (C, E). The data are presented as an average of at least three independent experiments. The graphs in panels (B), (C), (D) and (E) represent the data obtained at 10 μM dTTP and 50 nM PARP1 or PARP2. The data are presented as an average of at least three independent experiments and showed the mean values ± SD. (F) The reaction products of dTMP incorporation by Polβ (lanes 3–6) in the presence of PARP1 (left part) or PARP2 (right part, lanes 7–10) when outward- (upper panel) or midward-oriented (lower panel) 5′-FAM–labelled gap-NCP was employed. Lane 1: substrate AP-NCP, lane 2: substrate gap-NCP. In all cases, Polβ concentrations of 2.5 and 50 nM were chosen as the reaction conditions for substrates out-NCP and mid-NCP, respectively. The reaction products were separated on a 15% denaturing polyacrylamide gel.

In all the cases, the maximal velocities of dTMP incorporation were dramatically lower in the presence of PARP1, down to total inhibition of DNA synthesis at the PARP1 concentration of ≥100 nM. Additionally, an increase in *K*_M_ values for out-NCP was observed [*K*_M_ (out-NCP) ≈ 20.5 ± 1.7 μM] in the presence of 100 nM PARP1. The PARylation catalysed by PARP1 reverted of both kinetic parameters but did not reverse them to their initial levels (Figs. 4A, 4C, 4E and S3).

PARylation by PARP1 was carried out to restore the Polβ polymerase activity; however, in all cases, the reaction yield did not reach to the initial level (Fig. 4C). It is noteworthy that the bell-shaped peak of the curves for substrate mid-NCP slightly shifted to the side where higher concentrations of NAD^+^ were applied.

An important component of the BER processes is XRCC1. Its role is largely described as a scaffold protein for other BER participants (48). There are a lot of data concerning the modulation of Polβ activity by XRCC1 on naked DNA and in the nucleosomal context. In our study, the influence of PARPs and PARylation on Polβ activity was investigated in the presence of XRCC1 too.

Basically, in the assay of Polβ activity, the presence of XRCC1 did not lead to dramatic changes in the kinetic parameters of DNA synthesis at the concentrations close to Polβ levels (data not shown). This effect was independent from the substrate type.

Nevertheless, XRCC1 had a slight protective effect against PARP1-driven inhibition of Polβ activity, where the influence of XRCC1 in the assay with out-NCP was more significant (Figs. S3, 3C, and 3D).

The presence of XRCC1 during the PARylation catalysed by PARP1 did not cause a significant change in DNA synthesis efficacy regardless of the NCP substrate type (Fig. 4C and 4D). Moreover, the shape of the curves reflecting the efficacy of dTMP incorporation under PARylation in the presence of XRCC1 was the same as the shapes seen in the absence of XRCC1 (Fig. 4C and 4D).

The presence of PARP2 also as PARP1 decreased the Polβ activity (Figs. 4E, 4F and S3B, S3D). Indeed, it decreased the maximal velocities of dTMP incorporation in all cases [*k*_cat_ (out-NCP) ≈ 26.7 ± 2.5 10^−3^ s^−1^ and *k*_cat_ (mid-NCP) ≈ 5.1 ± 0.6 10^−3^ s^−1^] but not as dramatically as PARP1 did for substrate mid-NCP. An increase in *K*_M_ values in the presence of 100 nM PARP2 was observed too [*K*_M_ (out-NCP) ≈ 21.2 ± 1.8 μM and *K*_M_ (mid-NCP) ≈ 5.7 ± 0.7 μM]. Therefore, PARP2 caused inhibition of the DNA synthesis catalysed by Polβ.

The PARylation catalysed by PARP2 did not positively affect the intensity of DNA synthesis in the wide range of NAD^+^ concentrations, especially for substrate out-NCP (Fig. 4E).

The main difference of the effects of PARP2 and PARylation from the effects of PARP1 on Polβ activity was attributed to the influence of XRCC1. The presence of the scaffold protein substantially weakened the PARP2 inhibition on both NCP structures (Figs. S3B, S3D, 4E and 4F). Moreover, an additional decrease in the dTMP incorporation level was observed at the highest concentrations of NAD^+^ (Fig. 4F).

### The LigIIIα activity and the effects of PARP1 and PARP2

The last stage of the repair process is nick sealing. The main role at this step of BER is ascribed to LigIIIα. Accordingly, a lot of data support specific complex formation between DNA LigIIIα and XRCC1; the DNA ligase activity on an NCP has been investigated in the absence or presence of equimolar concentration of XRCC1 (49, 50). In our study, well-quantifiable data were not obtained with a mid-NCP substrate in the reaction with LigIIIα. Odell and co-authors have reported that the efficiency of LigIIIα in complex with XRCC1 is unaffected by the helical orientation of the nick in nucleosomes (51). Besides, those authors supposed that the complex of LigIIIα with XRCC1 can disrupt nick-containing nucleosomes. The model NCP tested in our study contains mid-ward oriented lesion at the seventh position from the 5′ end of a DNA duplex; consequently, it could be unstable and dissociate from the NCP under LigIIIα action, consistently with the above suggestion. For this reason, further experiments were performed on substrate out-NCP only.

It was determined that the use of a higher concentration of LigIIIα does not dramatically change the level of the final 5′-FAM–labelled reaction product but drives the formation of a protein–nucleic acid complex that did not resolve during electrophoresis (Fig. S5). To decompose these complexes, we conducted treatment with Proteinase K in all types of the reactions and with PARG in the reactions with NAD^+^. In addition, we switched to a more sensitive label, isotope 32 of phosphorus, present at one of the 5′ end of a DNA duplex of NCP (Figs. 5 and S4). It should be noted that a small amount of the full-length DNA strand was detectable in these protein–nucleic acid complexes but not the short part of the DNA substrate that is marked as a primer in the figures. Therefore, the extent of ligation was evaluated by means of the proportion of the unreacted primer.

**Figure 5.**
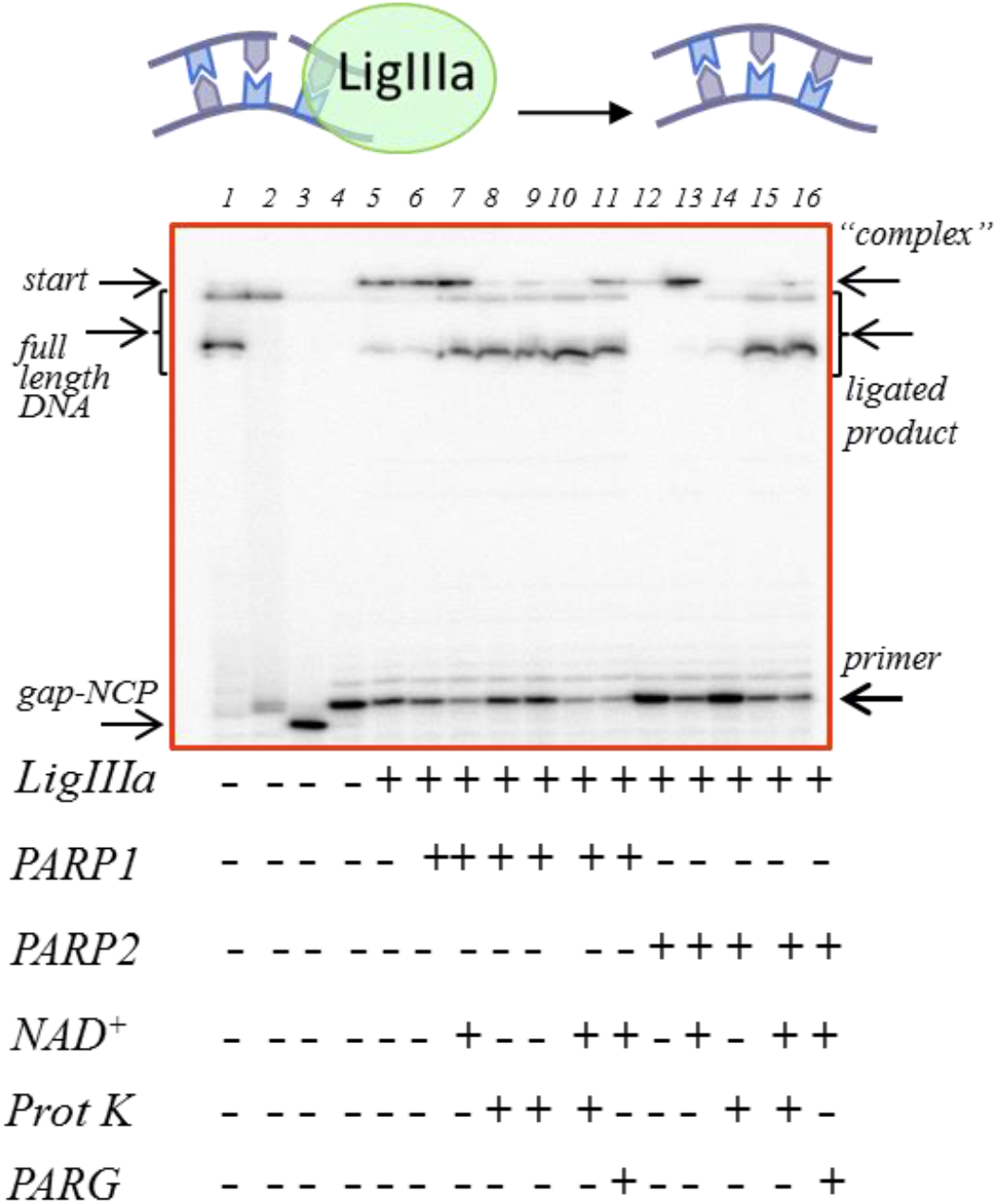
The LigIIIα activity on nicked substrate out-NCP. Product separation after the sealing of 5′[^32^P]labelled nicked out-NCP by LigIIIα with PARP1 or PARP2 and PARylation in the presence of XRCC1. Lane 1: native out-NCP; lane 2: out-NCP incubated with UDG; lane 3: out-NCP incubated with UDG and APE; lane 4: out-NCP incubated with UDG, APE, Polβ and dTTP, resulting in nicked out-NCP; lanes 5–16: nick sealing in the presence of LigIIIα (lanes 5, 8) and PARP1 (lanes 6, 9) or PARP2 (lanes 12, 14) without or with NAD^+^ (lanes 7, 10, 11 and 13, 15, 16, respectively). Lanes 8–10 and 14–15 correspond to lanes 5–7 and 12–13 with an additional treatment (with Proteinase K), respectively. Lanes 11 and 16 correspond to lanes 7 and 13 with an additional treatment (with PARG). The data are presented for experiments involving 20 nM NCP, 500 nM LigIIIα, 500 nM XRCC1, 100 nM PARP1 or PARP2 and 100 μM NAD^+^.

The influence of PARP1 and PARP2 on the LigIIIα activity was investigated in the absence or presence of NAD^+^ (Fig. 6). The following conclusions can be drawn from these experiments. PARP1 as well as PARP2 exerted an inhibitory effect on the LigIIIα activity under the experimental conditions (compare the columns of the first group with the second and fourth one in Fig. 6). PARylation attenuates their inhibition and increases the yield of the reaction product. It should be noted that PARP2 exerts these effects to a greater extent than PARP1 does (compare the difference in bars between the second and third group with the difference between the fourth and fifth group in Fig. 6). The presence of XRCC1 had no dramatic impact on the LigIIIα activity in our model system (bars of the first, third, fourth and fifth groups in Fig. 6); however, it slightly attenuated the sealing inhibition caused by PARP1 (bars of the second group in Fig. 6).

**Figure 6.**
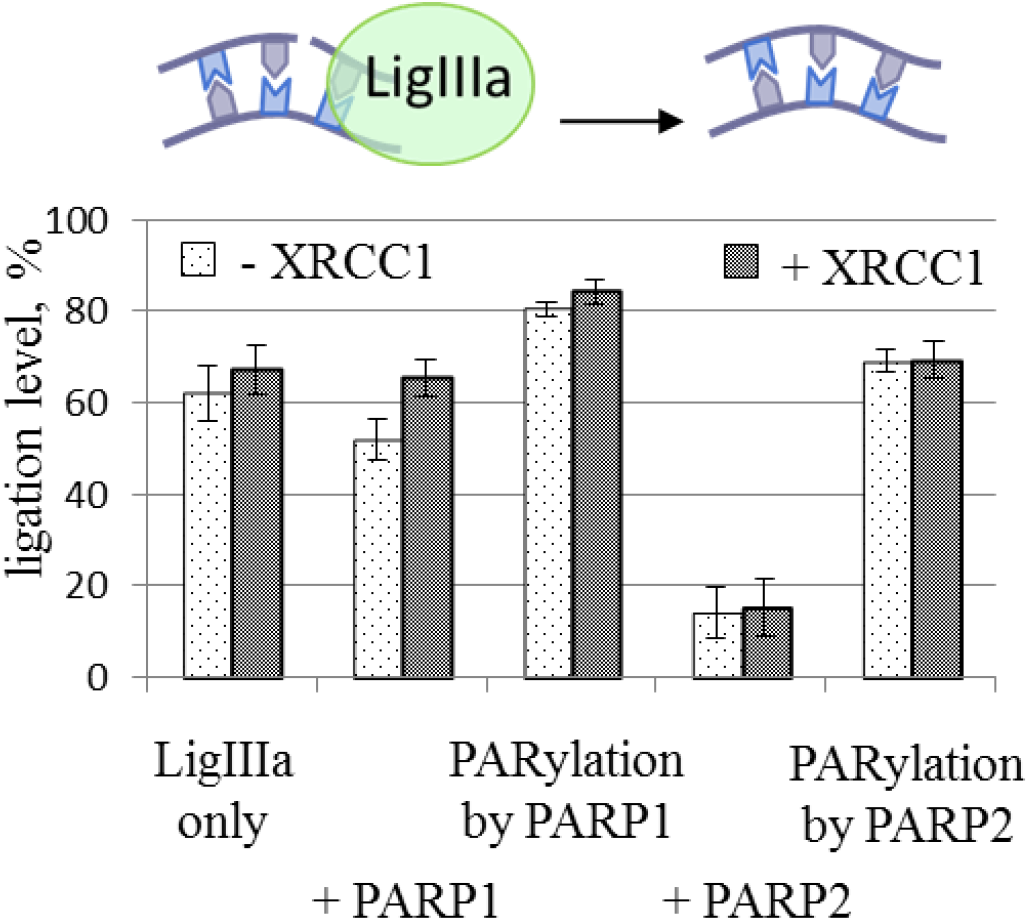
The influence of PARP1 or PARP2 and PARylation on the LigIIIα activity in the absence or presence of XRCC1. The ligation magnitude of the 5′[^32^P]labelled nicked NCP substrate under the action of LigIIIα was affected by PARP1 or PARP2 and PARylation in the absence or presence of XRCC1. The ligation magnitude was calculated as a percentage of the unreacted primer subtracted from 100%. The data are provided from experiments involving 20 nM NCP, 500 nM LigIIIα, 100 nM PARP1 or PARP2 and 100 μM NAD^+^. The data are presented as an average of at least three independent experiments and showed the mean values ± SD.

In summary, our experiments point to a specific influence of PARP1, PARP2 and PARylation on the activity of the main BER enzymes (APE1, Polβ and LigIIIα) with XRCC1 on an NCP. It was found that PARP1 suppresses the activity of APE1, Polβ and to a smaller extent LigIIIα. PARP2 exerts a significant effect on the activity of LigIIIα only. The decreases in the activities of APE1 and Polβ are reversed in the presence of the PARylation catalysed by PARP1 and to a lesser extent by PARP2. Meanwhile, the complementation with XRCC1 protects the dNMP transferase activity of Polβ from the inhibition by PARP1 and PARP2. Moreover, this complementation slightly affects the intensity of DNA synthesis under the PARylation catalysed by PARP1 and leads to a decrease of Polβ activity in case of PARylation by PARP2, in a NAD^+^-dependent manner. The remarkable result was obtained in the assay of DNA sealing by LigIIIα. This reaction was stimulated by the PARylation catalysed by PARP2. It is worth mentioning that outward-oriented lesions are more accessible to all the enzymes, and their processing requires a lower enzyme concentration and PAR level. A higher level of PAR is needed for the repair of more hidden damage sites.

## DISCUSSION

At present, a substantial amount of data about the interaction of PARP1 or PARP2 with various DNA structures is available (8, 30-32, 45, 52-55, 56, 57). The data on the affinity of PARP1 and PARP2 for structures are quite different among these works. The information about the structure of DNA substrates effective at PARP1 or PARP2 activation is also quite different among various sources (8, 30, 45). On the one hand, this fact can be explained by the dissimilar approaches applied in the studies that often require various experimental conditions. On the other hand, the reason for the discrepancies may be the use of dissimilar DNA models. For example, during interaction with blunt ends of approximate 30-40 bp DNA duplexes, PARP1 or PARP2 can come into contact with a nearby internal damage site. In the study (55), AFM analyses were performed on double-strand DNAs more than 1 kbp long containing damage at a substantial distance from the blunt ends and thus excluding an overlap of PARP1 or PARP2 interaction with various damage sites. It was found that PARP1 manifests high affinity for a single-strand break in DNA and similar affinity for blunt ends; PARP2 showed high affinity for a single-strand break and low affinity for blunt ends. The activity of both proteins toward the analysed DNA with a given type of damage correlated with these proteins’ affinity, though the activity of PARP2 was always lower as compared to PARP1. Additionally, in search of a possible specific role of PARP1 and PARP2, the interactions of these proteins with DNA structures mimicking various stages BER intermediates have been investigated by AFM (31). In that study, the AFM data indicated that PARP1 has higher affinity for early BER intermediates containing an AP site or a gap with a 5′-deoxyribose phosphate than PARP2 does, whereas PARP2 interacts more efficiently with a 5′-phosphorylated nick and may contribute to the regulation of the final ligation step.

In our work, we determined the contribution of different DNA lesions such as blunt end, AP site or 5’-phosphorylated gap to PARP1 or PARP2 interaction with components of an NCP (Table 1). As for PARP1, it did not exert significant difference for any substrates independent of the existence of damage site or nucleosomal structure that could be explained by high affinity of the protein for blunt-ended part of DNA duplex. This observation is in good agreement with previously reported data on PARP1 affinity (55).

The behaviour of PARP2 under the interaction with the substrates was more complicated (Table 1). PARP2 manifested extremely high affinity for gapped DNA that is in line with previously published findings (31, 55). Additionally strong increase of PARP2 affinity for gapped NCP may suggest a significant contribution of PARP2’s interaction with the histone core of an NCP.

EMSA results point to the complexity of the interaction of each PARP with different NCPs owing to the presence of several protein–nucleic acid complexes with different mobility, which could correspond to independent and simultaneous formation of complexes between PARP and blunt ends of DNA, a damage site, and/or histones (Fig. S2H and S2I). Overall, based on our results we can suggest that during the association of PARP1 with an NCP, the protein–nucleic acid interactions may be predominant. During the association of PARP2 with a damaged NCP two different processes could compete with each other: protein–protein and protein–nucleic acid interactions.

It is noteworthy that the activation of PARP1 upon interaction with DNA in any cases poorly correlates with the enzyme’s affinity for DNAs and is much higher on nick-containing duplexes (52, 54). This fact can be explained by the specific conformational changes of PARP1 structure in the complex with damaged DNA upon NAD^+^ binding and after activation (58-60). Furthermore, PARP1 exerts an additional specificity under interaction with NCP. The spFRET microscopy experiments has indicated that *in vitro* binding of PARP1 to chromatin leads to a considerable change in nucleosome structure through a significant increase in the distance between the adjacent gyres of nucleosomal DNA; this phenomenon may be described as nucleosome unfolding and can be almost completely reversed by autoPARylation (61). In this study, we investigated the PARPs’ activity under their interaction with native or damaged NCP by real-time measurement of fluorescence anisotropy as well as by the detection of the total amount of synthesised PAR.

It was obtained that the *k*_obs_ values after PARP1 PARylation were dependent on the damage type and were higher for gap-NCP at the comparatively equal *K*_M_ values (Table 2). This variation may reflect the activation efficacy of PARP1 in the presence of undamaged DNA or various types of DNA damage: the AP site or gap (25, 31, 55). The existence of a specific damage site in the DNA in addition to a blunt end could result in binding of additional PARP1 molecules to the lesion. In turn, this event leads to modulation in the efficiency of PAR synthesis and thereby to changing in the rate of dissociation of autoPARylated PARP1 from the complex with a damaged NCP characterised by higher *k*_obs_ values.

Under evaluation of the impact of PARylation efficiency using our nucleosomal structures, we found that PARP1 is the most efficiently activated upon the interaction with native or AP-NCP, but not with gapped NCP (Fig. 2, A-D). It is possible that the presence of a lesion in the nucleosomal DNA can significantly affect the geometry of the PARP1-NCP complex. In our structure, the lesion is located close enough to the blunt-ended part of the DNA helix (less than one and less than one and a half turns of the DNA helix for midward and outward-oriented damage, respectively). Possibly, such a mutual arrangement of two PARP1 binding sites leads to the formation of a nonproductive PARP1-NCP complex where more than one PARP1 molecule binds to the substrate, which corresponding alternates the amount of PAR. At the same time, an increasing of *k*_obs_ values upon introducing a lesion to the DNA part of the nucleosome, at the absence of a significant difference in the PARP1 affinity for the substrate, could be consequences of the dissociation of a less productive complex that requires a smaller amount of ADP-ribose polymer.

PARP2 similarly does not manifest a direct correlation between the affinity for a DNA structure and activation efficiency (8). Recent studies, however, including studies on nucleosomes, revealed certain features of DNA structures such as various DNA breaks with a 5′-phosphate group that necessary for the efficient binding and/or activation of PARP2 (30-32, 45, 55, 57). In our experiments, all the analysed *K*_M_ and *k*_obs_ parameters were highest for the gapped substrate (Table 2). At the same time, the efficiency of PARP2 activation by different NCP substrates obeys the same behaviour that of PARP1 - the least amount of ADP-ribose is synthesized in the presence of gapped NCP (Fig. 2, I-H). Nevertheless, the affinity of PARP2 for this substrate is a one order of magnitude higher than for NCP with an intact or AP-containing sugar phosphate backbone (Table 1). The efficacy of PARP2 activation can be explained by the formation of a stronger, but less productive PARP2-gapped NCP complex. Under investigation of the PARylation by the fluorescence anisotropy at a low NAD^+^ concentration, as indicated by a part of the titration curve with a small slope up to 200 μM NAD^+^, a larger contribution to the *k*_obs_ constant makes the interaction of PARP2 with NAD^+^ molecule (Fig. S2E and S2G). At a high NAD^+^ concentration, the *k*_obs_ constant to a greater extent reflects the dissociation rate of the complex, as evidenced by the increasing amplitude of *k*_obs_ curves with increasing NAD^+^ concentration above 1250 μM. Thus, it is possible that the dissociation of PARylated PARP2 from the complex with gapped NCP takes longer but derives more efficient under increase of NAD^+^.

It should be noted that the amount of PAR synthesized by PARP2 upon the activation by different types of naked DNA does not vary by the structures, although the affinity of PARP2 for gapped DNA is also significantly higher. Additionally, PARP2 generally proved to have low affinity for a DNA compare to a NCP. It is possible that in the absence of histones and/or a more rigid nucleosomal structure upon the interaction with DNA, the PARP2-gapped DNA complex exhibits an excessive conformational dynamic, which makes it possible to increase the amount of the productive complex and, accordingly, to increase the PAR yield.

Therefore, collectively our data are suggestive of the expected contribution of PARP1 starts from the initial stages of BER. As for PARP2, its contribution is expected at later stages of the BER process, starting from the formation of gapped substrates. The character of the interaction of PARP2 with the damaged NCP reveals a significant impact to the interaction with nucleosomal proteins.

APE1 is the main enzyme for an AP site repair, which at the earlier stage of BER cleaves the sugar-phosphate backbone with a formation of 3’-hydroxyl and 5’-dRp groups. Its activity is highly dependent on the bases surrounding an AP site and on secondary structure of the DNA substrate (reviewed in 62, 63). In the case of an NCP, each nucleotide has a specific orientation relative to the histone surface; this orientation may make the nucleotide either more or less accessible to a repair enzyme depending on the mechanism of the interaction. It turns, the orientation of the AP site towards the histone core is important to realisation of APE1 activity (64-66). So, in our experiments we used the specific conditions to ensured equivalent APE1 efficiency using an midward- or outward-oriented AP site containing NCP.

We observed the suppression of the APE1 activity in the presence of PARP1 and substantially restoration of the AP site cleavage level at the PARylation regardless of the damage orientation. These results are consistent with previous studies on model naked DNA (25, 47). These observations can be ascribed to the competition between APE1 and PARP1 for the binding to an AP site. The interaction of PARP1 with an AP site has been demonstrated previously, and at high concentrations, PARP1 can compete with APE1 for the binding to an AP site (25). In addition, PARP1 forms complexes with the blunt-ended part of DNA and thus can sterically prevent the interaction of APE1 with the closely located AP site. The PARylation initiated by the binding to damaged DNA results in the formation of a negatively charged polymer on the PARP1 surface thereby promoting its dissociation from DNA due to electrostatic repulsion and opens the access of APE1 to the substrate. Therefore, the PARylation catalysed by PARP1 restored the AP endonuclease activity of APE1 and diminished its inhibition by PARP1.

PARP2 slightly affected APE1 activity regardless of substrate complexity (naked DNA or NCP), and PARylation did not reverse this inhibition. This finding can be explained by the low affinity of PARP2 for AP site–containing DNAs revealed in this study and earlier as well as by the comparatively low reaction rates of PARP2 under PARylation (31, 47). On the other hand, PARP2 showed stronger affinity for NCP than for naked DNA. Taken together, these data further suggest the notion that PARP2 interacts with an NCP not only through the DNA component but also through the histones and likely their PARylation or other posttranslational modifications can influence on PARP2 interaction with gapped NCP.

The formation of the gap with 3’-hydroxyl and 5’-dRp groups in a DNA requires the DNA polymerisation activity for BER extension, specifically the activity of Polβ. This enzyme form the most stable complex with scaffold BER protein XRCC1 among the complexes of the other key BER proteins with XRCC1 (27). Some investigators discuss the contribution of the XRCC1–LigIIIα complex to the interaction with Polβ (67). Mostly, the literature data support the coordinating role of XRCC1, which has special regions for binding to LigIIIα and Polβ (28, 68–70). A recent study uncovered the formation of a stable ternary complex, XRCC1–Polβ–LigIIIα (71). The authors of ref. (51) propose a molecular mechanism based on the disruption of nucleosomal structure by the complex of XRCC1 and LigIIIα, thereby resulting in easier access of Polβ to the gap. In our experiments we tested the activity of Polβ to carry out the dNMP incorporation in the absence or presence of the XRCC1 and the influence of PARP1/PARP2 and PARylation on this activity using in- and out-NCP.

It is possible that the inhibition of DNA synthesis by PARP1 and the subsequent increase in the Polβ reaction yield in the presence of NAD^+^ is a consequence of the following effects. It should be noted that the binding of PARP1 to a nucleosome causes partial NCP de-compaction in the vicinity of a double-strand break (61). Such de-compaction may increase the access of Polβ to the damage site and enhance the enzymatic activity towards our NCP models. Moreover, an increase in Polβ activity after damage dislocation towards the blunt ends has been documented (72). Nevertheless, the proximity of the single-strand break to the blunt ends of DNA duplexes may start protein–protein competition for the substrate between PARP1 and Polβ. Under PARylation conditions, the inhibition is attenuated by the dissociation of PARylated PARP1. The recovery of Polβ activity in the presence of PARP1 PARylation has been demonstrated by means of various model DNAs mimicking BER intermediates (6, reviewed in 26).

Of note, the dependence of the polymerase reaction yields on the NAD^+^ concentration has a bell-shaped profile. Meanwhile, the inhibition of DNA synthesis at a high concentration of NAD^+^ could be explained by the dissociation of the ternary complex NCP–PARP1–Polβ owing to total protein PARylation (Fig. S4). The PARylation of Polβ as well as its interaction with PAR were recently reported (73 and on Fig. S4). The curve bias seen in Fig. 4C plausibly is a consequence of impeded access to the damage site, necessitating a higher PAR level in the presence of Polβ; therefore, it requires higher concentrations of PARP1 and NAD^+^.

Owing to specific complex formation between Polβ and XRCC1, the competition for the DNA substrate between the DNA polymerase and PARP1 in the presence of the scaffold protein is not so critical; this situation allows to attenuate the inhibition of Polβ activity by PARP1. In this case, under PARylation conditions, the same concentration of NAD^+^ was needed for the recovery of Polβ activity in the absence of XRCC1 possibly because the presence of XRCC1 means an additional direct PARylation target (Fig. S4) (33).

Supposedly, the interaction of PARP2 with the gapped NCP is different from such an interaction of PARP1 and may be mostly promoted by the interaction with the gap (Table 1). Moreover, the interactions of PARP1 and PARP2 with gapped model DNAs have been demonstrated earlier (22, 23, 47). Given that PARP2 has comparatively low affinity for blunt ends, it has only a slight steric effect on Polβ functioning during the interaction with an NCP – in comparison with PARP1 – and largely competes with the polymerase at the gap site. Besides, PARP2 has lower efficacy of PARylation as compared to PARP1 (47, 74). Consequently, the amount of PAR in this case is not enough for dissociation of PARP2 from the complex with gap-NCP and for promoting Polβ activity. It is likely that the addition of XRCC1 results in the formation of its specific complex with Polβ; this complex can efficiently compete with PARP2 for the substrate binding, thereby attenuating PARP2’s inhibitory effect on DNA polymerase activity. At the same time, XRCC1 is a good target for PARylation by PARP2, and at a high NAD^+^ concentration, the PARylation could induce the dissociation of the active proteins from the NCP; this phenomenon may explain the reduction in polymerase activity (Fig. S4).

At the last stage of BER DNA LigIIIα in the complex with XRCC1 is sealed the single-strand break to restore the integrity of DNA. In our study we investigated the influence of PARP1/PARP2 and PARylation on the ligation and the impact of XRCC1 to the reaction.

It is possible that a strong effect of XRCC1 on the activity of LigIIIα was not observed because of the damage localisation in a nucleosome region having greater flexibility (75, 76). We found that the presence of XRCC1 slightly attenuated the sealing inhibition caused by PARP1. This result is consistent with the published data on the inhibitory effect of XRCC1 on the PARP1 activity (77). Under the assumption that LigIIIα binds to a nicked NCP and partially disrupts it, the inhibition of its activity by PARP1 could be explained by the competition between the proteins upon substrate binding close to blunt ends of the DNA duplex. Aside from the competitive interplay, the interaction of PARP2 with the nicked NCP probably prevents the nucleosome disruption by LigIIIα and as a consequence causes a lower reaction yield.

The increase in the sealed product level in the presence of NAD^+^ could be ascribed to the following scenario. In this case, not only a PARP molecule but also XRCC1 and probably LigIIIα may undergo modification. It has been shown that LigIIIα can detect DNA nicks in the presence of either PAR or PARylated PARP1 (78, summarised in 15). Moreover, the LigIIIα zinc finger, which – just as the catalytic domain – is important for the detection of damaged DNA, may facilitate recognition of the DNA nick in the presence of the negatively charged PAR (78, 79). Moreover, Caldecott’s group has discovered that XRCC1 can be attracted to a damage site through the interaction with PAR, and the catalytic activity of either PARP1 or PARP2 can promote the loading of endogenous XRCC1 onto the damage site, possibly causing the formation of the specific binary XRCC1–LigIIIα complex at this site (80, 81). Hence, a certain PAR level will stimulate the formation of the LigIIIα specific complex at the DNA damage site. With a further increase of the PAR level in the system, LigIIIα may be pulled down from the NCP because of the competition between PAR and DNA as binding substrates or a breakdown of the entire complex owing to the formation of a large amount of the negatively charged polymer of ADP-ribose. In this case specific interaction and/or activity of PARP2 with a NCP could provide more accurate adjustments for the LigIIIα or XRCC1–LigIIIα activity and stimulate ligation.

## CONCLUSIONS

It is clear that nuclear PARPs trigger and control DNA repair under genotoxic stress, and PARylation regulates these processes. PARP1 generates the signal of PARylation and interacts with histone and non-histone chromatin proteins and XRCC1 through the PAR (82-84). Protein–protein interactions of PARP1 with the other BER proteins have been estimated quantitatively in the absence or presence of different BER DNA intermediates, and their contributions to the BER process were evidently supported by the data (27, 28). Efficient interaction of PARP1 with APE1 as well as with Polβ has been demonstrated. In turn, XRCC1 interacts with several BER enzymes: (i) APE1 via a linker and the BRCT1 domain (48); (ii) Polβ via the N-terminal domain (68); and (iii) LigIIIα via the BRCT2 domain (85). XRCC1 also interacts with PARP1 via the BRCT domain and with PARP2 through the WGR domain but is recruited mostly by recognising PAR formed near DNA damage (29, 33, 86). In the same way, PARP2 interacts with the main BER proteins, i.e. PARP1, XRCC1, Polβ and LigIIIα, as well as with the damaged DNA; accordingly, it is possible that the WGR domain of PARP2 participates in the protein–protein and nucleic-acid–protein interactions (29, 45, 56). On the other hand, the interactions of PARP2 with the BER proteins have not been estimated quantitatively. Furthermore, the data on the affinity and activation of PARP1 and PARP2 towards damaged DNA indicate that PARP1 prefers earlier BER-intermediates, whereas PARP2 mostly acts on later ones (30, 31, 87). In addition, this study as well as literature data indicate functional dependence of various repair enzymes’ activities on PARylation and PAR synthesis (reviewed in 26). According to these findings, the influence of PARP1 on BER processes should manifest itself before and at the stage of APE1-induced cleavage and gap/break formation. PARP2 may be more important at the ligation stage because of both the competitive interaction with LigIIIα during DNA binding and during the ligase activation by PAR production. On the basis of the results presented here and in the literature, the following hypothetical model of the regulation of basal BER process in the absence of massive DNA damaging exposure, by PARP1 and PARP2 activities can be formulated.

The occurrence of the DNA damage triggers the activation of a signalling pathway through PARP1. In this process, some regions of the compacted chromatin could be converted to a partially de-compacted state. PARylation enables the dissociation of histone and non-histone chromatin proteins and involves specific repair proteins (88). The main role of PAR at this stage is to loosen chromatin structure and to attract BER proteins such as scaffold protein XRCC1 and possibly some crucial proteins such as APE1 and others. The interactions of XRCC1, APE1 and other BER proteins with PAR have been documented (73, 77, 89). Co-localisation of PARP1 and APE1 on an intact or incised AP site has been detected by AFM, meaning that the direct interaction of PARP1 and APE1 may facilitate the search for DNA damage sites (90). The cleavage of an AP site by APE1 leads to additional activation of PARP1 and PARylation resulting finally in dissociation of PARP1. It makes the way for subsequent stimulation of the BER process by the removal of APE1 from its product by Polβ, followed by a dRP-lyase reaction and repair synthesis (91). The weight of evidence suggests that the functions of PARP1 and PARP2 probably partially overlap after the AP site cleavage and supports a role of PARP2 in the regulation of BER activity. Both PARP1 and PARP2 engage in a specific interaction with the product of AP site cleavage, and both PARPs are activated by gap substrates (23, 65). Nonetheless, we believe that the contribution of PARP2 to BER becomes more specific from this point forward and begins to increase. The presence of a specific single-strand break DNA structure as well as PAR synthesised by PARP1 attracts PARP2, which in turn synthesises additional PAR molecules. PAR can regulate the turnover of BER proteins at various stages of the process. Whereas PARP1 and PARP2 regulate Polβ activity, PARP2 stimulates LigIIIα activity to complete the repair process, indirectly through their interactions with XRCC1 and PAR. The synthesis of PAR that is catalysed by PARP2 during the interaction with damaged DNA helps the Polβ–XRCC1 complex to complete its action and stimulates the activity of LigIIIα–XRCC1 towards the NCP. The PAR level leads to a breakdown of the specific ligase complex, and repair is thus accomplished. We propose that the PAR synthesised at the initial stages by PARP1, and at the final steps also by PARP2, can act as a regulatory factor for the step-by-step progression of the BER process.

## MATERIALS AND METHODS

### Materials

Synthetic oligonucleotides, including 5′-FAM–labelled oligonucleotides, were acquired from Biosset (Novosibirsk, Russia). Reagents for electrophoresis and basic components of buffers were purchased from Sigma (USA). γ[^32^P]ATP and α[^32^P]ATP (with specific activity of 5000 and 3000 Ci/mmol, respectively) were bought from the Laboratory of Biotechnology (Institute of Chemical Biology and Fundamental Medicine, Novosibirsk, Russia), whereas recombinant T4 polynucleotide kinase and *E. coli* uracil-DNA glycosylase from Biosan (Novosibirsk, Russia). Proteinase K was bought from NEB (USA), and ultrapure dNTPs, ddNTPs, NTPs and NAD^+^ from Promega (USA). The synthesis of radioactive NAD^+^ was carried out according to ref. (92).

### Protein purification

Human LigIIIα and XRCC1 were purified according to refs. (71, 93), whereas human APE1, rat Polβ and PARG according to (94–96). Human PARP1 and mouse PARP2 were purified as described elsewhere (97).

### Oligonucleotide substrates

Oligodeoxyribonucleotides were 5′[^32^P]phosphorylated by T4 polynucleotide kinase as described before (92). The amplification of 5′-FAM- or ^32^P-labelled DNA and subsequent NCP reconstitution were performed according to (98). The purity of the sample was estimated by the homogeneity controlled by electrophoretic mobility under on a 10% polyacrylamide gel under non-denaturing conditions. The assembly of the nucleosome was considered correct when one band was detected on the electropherogram with electrophoretic mobility corresponding to NCP one (Fig. S1). On the basis of molecular modelling and biochemical data about the nucleobase orientation of the 601 Widom sequence, two positions of the 603 Widom sequence with different damage orientations were chosen; they were either outward- or midward-oriented lesions (99, 100).

The following sequences of primers were used for the assembly of a DNA and NCP with a middleward- or outward-orientation of the damage site: 5′-CCCAGT<u>U</u>CGCGCGCCCACC and 5′-ACCCCAGGGACTTGAAGTAATAAGG for the substrate with a midward-oriented lesion, and 5′-ACCCCAGGGACT<u>U</u>GAAGTAATAAGG and 5′-GGGTCAAGCGCGCGGGTGG for the substrate with an outward-oriented lesion (Fig. 1B). At the first stage, specific conditions for AP site generation were found that ensured equivalent UDG efficiency using a midward- or outward-oriented NCP and DNA (101). In all cases, the AP sites were generated by the UDG activity during direct incubation of a U-containing DNA or U-NCP solution with the enzyme for 30 min at 37°C at the ratio of 1 activity unit per 0.6 pmol of DNA. The extent of reaction was controlled by the alkaline hydrolysis at the addition of 0.1M NaCl for 1 min at 37°C followed by probe heating for 5 min at 97°C and subsequent separation by electrophoresis on a 10% polyacrylamide gel under denaturing conditions. The produced AP site–containing substrates were subjected directly to the following reactions. Single-strand break substrates with a 5′-deoxyribose phosphate and a gap were prepared by incubation of 1 μM (for substrates with an midward-oriented lesion) or 0.1 μM (for substrates with an outward-orientated lesion) APE1 with 0.1 μM AP site–containing DNA (AP-DNA) or AP site– containing NCP (AP-NCP) in reaction buffer containing 5 mM MgCl_2_ for 15 min at 37°C and then were used for subsequent analyses. Low-salt nucleosome assembly buffer consisting of 10 mM NaCl, 0.2 mM EDTA, 5 mM β-mercaptoethanol, 0.1% of NP-40, 10 mM Tris-HCl pH 7.5, and 0.25 mg/ml BSA served as reaction buffer.

### Evaluation of K_d_

Values of dissociation constant *K*_d_ were determined by two methods. Reaction mixtures (final volume of 10 μl) for an electrophoretic mobility shift assay (EMSA) containing 50 nM 5′[^32^P]labelled DNA or NCP were incubated for 15 min at 37°C in reaction buffer with 0.01, 0.05, 0.1, 0.5 or 1 μM PARP1 or -2. To separate free DNA or NCP and nucleic-acid–protein complexes, the samples were then subjected to electrophoresis at 4°C on 4% native polyacrylamide gels in 1× TBE (Tris-borate-EDTA) buffer for 1 h at 100 V. After that, the gels were subjected to autoradiography and/or phosphorimaging for quantitation using a Typhoon imaging system from GE Healthcare Life Sciences and were analysed in OriginPro 7.5 (Microcal Software). Alternatively, *K*_d_ values were estimated by fluorescence anisotropy measurements based on the detection of real-time PARPs’ activity according to ref. (46). Briefly, a reaction mixture containing 0.03 μM 5′-FAM–labelled native, AP- or gap-DNA or NCP and 1.6–400.0 nM PARP1 or 0.016–2.000 μM PARP2 in reaction buffer (50 mM NaCl, 50 mM Tris-HCl, pH 8.0, and 5 mM DTT) with 5 mM MgCl_2_ was prepared on ice in a 384-well plate and incubated at room temperature for 10 min. The fluorescent probes were excited at 482 nm (482-16 filter plus dichroic filter LP504), and the fluorescence intensities were detected at 530 nm (530-40 filter). Each measurement consisted of 50 flashes per well, and the resulting values of fluorescence were automatically averaged. The measurement was carried out in kinetic scan mode. The measurements in each well were done 10 times with intervals of 3 min. The average values were used for the final plot, and *K*_d_ values were calculated by means of the MARS Data Analysis software (BMG LABTECH).

### The measurement of PARP’s activity

Efficacy of PARPs under activation by different substrates was evaluated as the amount of an poly(ADP-ribose) synthesised by PARP1/PARP2 using [^32^P]labelled NAD^+^ as a precursor. The reaction (in 60 μl) was initiated by the addition of 50 nM PARP1 or 100 nM PARP2 to a solution of 50 nM 5′-FAM–labelled native, AP- or gapped DNA or NCP in reaction buffer with 5 mM MgCl_2_ and 0-4000 μM NAD^+^ containing 1 μM [^32^P]labelled NAD^+^. The reaction was carried out at 37°C for 0, 1, 3, 5, 10 and 20 min and stopped by placing the aliquots of reaction mixture on paper filters (Whatman-1) soaked with a 5% solution of trichloroacetic acid. The filters were washed four times by the 5% trichloroacetic acid, then by 90% ethanol, and air-dried. The filters were subjected to autoradiography for quantitation on the Typhoon imaging system (GE Healthcare Life Sciences) and the quantity of radiolabel included into the acid insoluble fraction was analysed using Quantity One software (Bio-Rad). The quantitative data were analysed using MS Excel 2010 and presented on histograms as the Mean ± SD.

### The kinetic parameters of PARylation

*K*_M_ and *k*_obs_ values of the PARylation catalysed by PARP1 or PARP2 were estimated by fluorescence anisotropy measurements as described previously (46). Briefly, the reaction was initiated by the addition of an aliquot of a NAD^+^ solution of various concentrations (20–5000 μM) to the mixture of 0.2 μM PARP1 or 1 μM PARP2 with 0.1 μM DNA or NCP in reaction buffer with 5 mM MgCl_2_. The obtained values of the anisotropy decrease were normalised and plotted to calculate the *K*_M_ and *k*_obs_ values in the MARS Data Analysis software (BMG LABTECH).

### The influence of PARP1, PARP2 PARylation on the cleavage activity of APE1

First, the concentration of APE1 for effective AP site cleavage in the context of DNA or NCP was found. To this end, the reaction (in 10 μl) was initiated by the addition of 0.001–1.000 μM APE1 to a solution of 0.1 μM 5′-FAM–labelled AP-substrate in reaction buffer with 5 mM MgCl_2_. The reaction was carried out at 37°C for 0, 5, 10 or 15 min and stopped by the addition of loading buffer consisting of 7 M urea and 50 mM EDTA. The products were separated by electrophoresis on a 10% polyacrylamide gel under denaturing conditions and subjected to autoradiography for quantitation on the Typhoon imaging system (GE Healthcare Life Sciences) and were analysed using Quantity One software (Bio-Rad). APE1 efficacy was evaluated as the amount of an AP site cleavage product relative to all forms of DNA in a lane, expressed as a percentage. To assess APE1 activity toward naked DNA, a DNA duplex containing an AP site at the 13th position from the 5′-labelled end was used in all experiments. To evaluate APE1 efficacy in the presence of a PARP, the concentration of the enzyme that yielded 40–50% substrate cleavage within 15 min was chosen. This concentration was found to be 0.03 μM APE1 for substrates with an outward-oriented lesion, 1 μM APE1 for substrates with an midward-oriented lesion and 0.001 μM APE1 for naked DNA. For further experiments, in the same experimental scheme, PARP1 or PARP2 concentration was varied from 0.01 to 1.00 μM. At the last stage, the experiments with each PARP concentration were conducted at 400 μM NAD^+^. All reaction products were separated by 10% polyacrylamide gel electrophoresis and analysed as described above.

### The impact of PARP1, PARP2 and PARylation on the DNA polymerase activity of Polβ

At the first stage, the concentration of Polβ for effective gap-filling in the context of an NCP was found. For this purpose, the reaction (in 10 μl) was started by the addition of 0.001–1.000 μM Polβ and 100 μM dTTP to a 0.1 μM solution of the gap-substrate in reaction buffer with 5 mM MgCl_2_. The reaction was allowed to proceed at 37°C for 1, 3, 5 or 10 min and was quenched by the addition of loading buffer consisting of 7 M urea with 50 mM EDTA. The products were separated by electrophoresis on a 15% polyacrylamide gel under denaturing conditions and subjected to autoradiography for quantitation using the Typhoon imaging system (GE Healthcare Life Sciences) and analysed in the Quantity One software (Bio-Rad). The efficacy of the DNA synthesis was evaluated as the amount of the dTMP incorporation product relative to the initial gap-containing NCP form, expressed as a percentage. To evaluate Polβ efficacy in the presence of a PARP, the interval of 3 min was chosen, then the concentration of the enzyme was adjusted to ensure a linear range of the kinetics for substrates with an outward- or midward-oriented damage; thus, 2.5 and 50 nM, respectively, were chosen as suitable concentrations of Polβ. For further analyses, the kinetic parameters of the dTMP incorporation were estimated in the absence or presence of 1, 5, 10, 25, 50, 100 or 500 nM PARP1 or 20, 50, 100 or 500 nM PARP2. Additionally, the magnitude of dTMP incorporation was estimated in the presence of 1, 10, 100 or 1000 μM NAD^+^ for each PARP concentration. The effect of XRCC1 on the activity of Polβ was estimated in the same experimental scheme. The concentration of XRCC1 was selected at the first stage of the experiment to ensure dTMP incorporation at the same level, and was found to be 4 and 50 nM for substrates with an outward- or midward-oriented lesion, respectively. All reaction products were separated by 15% polyacrylamide gel electrophoresis and analysed as described above.

### The influence of PARP1, PARP2 and PARylation on the LigIIIα activity

At the first step, the reactions were carried out at various concentrations of LigIIIα – 0.01, 0.1, 0.5 or 1 μM – without or with XRCC1 in a molar ratio of 1:1 to LigIIIα to find optimal reaction conditions. Thus, the 0.5 μM concentration of LigIIIα was chosen. Gap-NCP substrates were treated with Polβ and dTTP to obtain a LigIIIα substrate as described above. The reaction mixtures (final volume of 10 or 30 μl) containing 0.02 μM 5′[^32^P]- or 0.1 μM 5′-FAM–labelled NCP substrate and 1 mM ATP in reaction buffer with 10 mM MgCl_2_ and 100 mM NaCl were incubated for 30 min at 37°C with the mixtures of LigIIIα and 100 μM NAD^+^; LigIIIα and 0.01, 0.1 or 1 μM PARP1 or PARP2; or LigIIIα, 100 μM NAD^+^ and 0.01, 0.1 or 1 μM PARP1 or PARP2 in the absence or presence of XRCC1. All probes were assayed as described above. In addition, 5′-FAM–labelled NCP-probes were assayed by 12% Laemmli SDS-PAGE at the 40:1 ratio of acrylamide to bis-acrylamide, followed by Coomassie Brilliant Blue R-250 staining. When indicated, before the electrophoresis, aliquots of the reaction mixture (10 μl) were treated with 0.001 U of Proteinase K or 0.5 μM PARG for additional 30 min at 37°C.

## ACKNOWLEDGEMENTS

We would like to express our deep gratitude to Dr. Vasily M. Studitsky (Fox Chase Cancer Center, PA, USA; Lomonosov Moscow State University, Moscow, Russia) for providing the DNA nucleosomal construct and for valuable advice on the design of the NCP. We thank Drs. Nina A. Moor, Maria V. Sukhanova and Rashid O. Anarbaev for helpful comments and fruitful discussions during the drafting of the manuscript. The English language was corrected and certified by shevchuk-editing.com.

