## Supplementary material for "The contribution of PARP1, PARP2 and poly(ADP-ribosyl)ation to base excision repair in the nucleosomal context": Figure S1

**SUPPLEMENTARY INFORMATION**

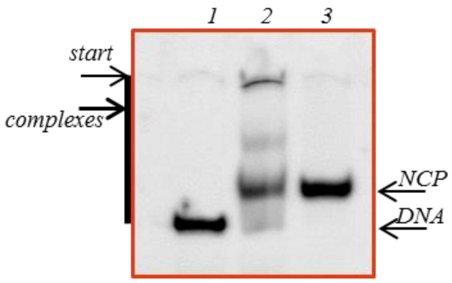


**Figure S1.** The electrophoretic mobility of 5'-FAM labelled products after the process of NCP reconstruction from model FAM-DNA and histone octamers in 4 % PAAG under non-denaturing conditions. Lane 1 — free DNA; lanes 2 — sample obtained by mixing of DNA and histone octamer with a ratio as 1:1 in low salt reaction buffer; lanes 3 — NCP assembled by gradient dialysis with the ratio of DNA and histone octamer with a ration as 1:1. NCP — nucleosome core particle, DNA — naked DNA, complexes – complexes of DNA with histone octamers.
