## Supplementary material for "The contribution of PARP1, PARP2 and poly(ADP-ribosyl)ation to base excision repair in the nucleosomal context": Figure S2

### SUPPLEMENTARY INFORMATION

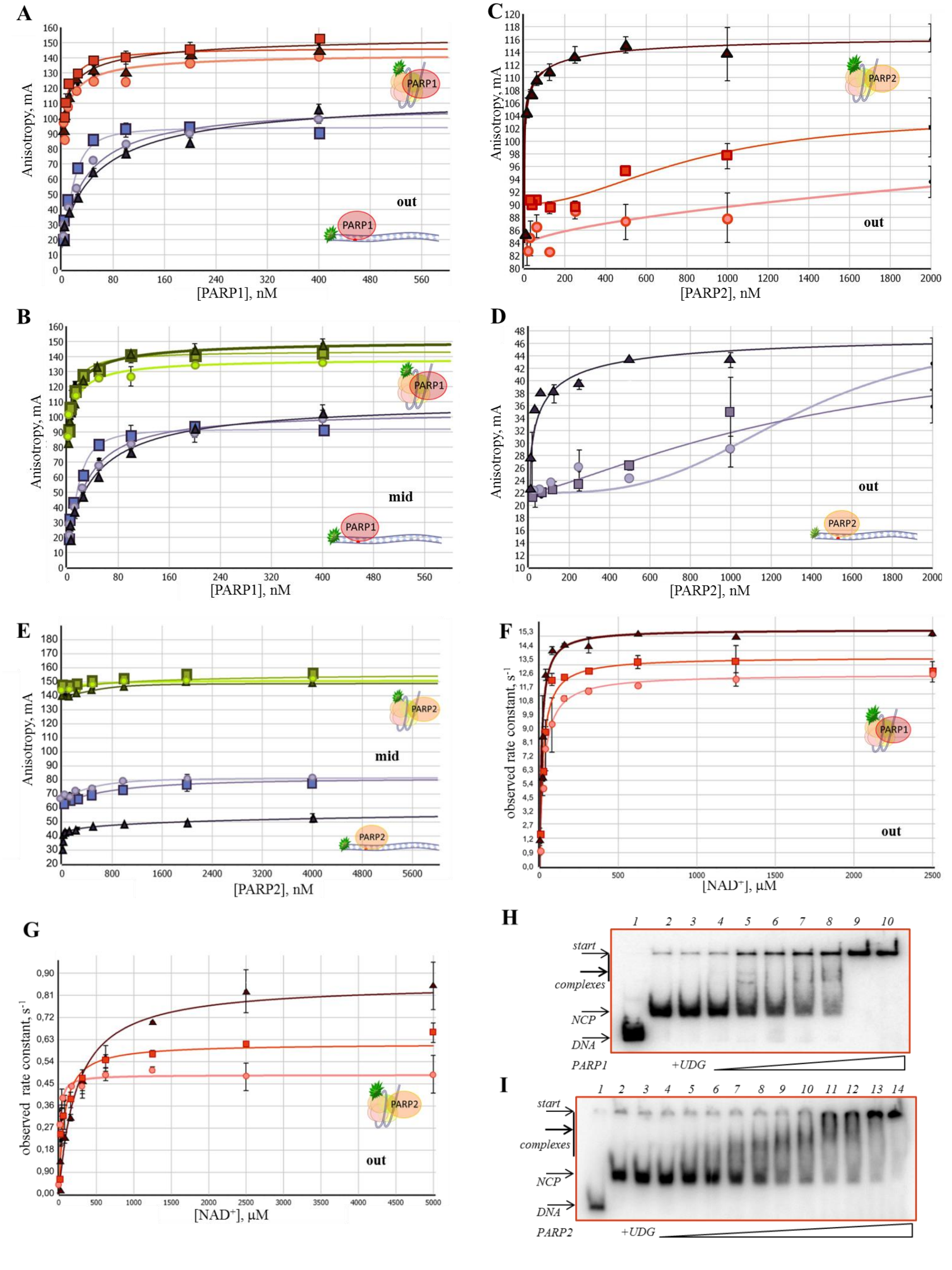

**Figure S2.** The affinity of PARP1 (A, B) and PARP2 (C, D, E) for native (circles), AP (squares) or gap-NCP (triangles) or DNA as measured by fluorescence anisotropy using 100 nM 5′-FAM–labelled NCP with an outward-oriented lesion (out-NCP; panels A, C, D) and the NCP with middle-oriented damage (mid-NCP; panels B, E). (F, G) Observed rate constant changes for protein–nucleic acid complexes of out-NCP with PARP1 (F) or PARP2 (G) under PARylation conditions. The reaction mixtures contained 30 nM 5′-FAM–labelled out-NCP, 100 nM PARP1 or 1000 nM PARP2 at various concentrations of $NAD^+$. In all the graphs, the experimental data on NCP substrates correspond to the red or yellowish-green curves; the DNA substrates are characterised by black, reddish-violet or violet curves. The data are presented as an average of at least three independent experiments. (H) EMSA analysis of the protein–nucleic acid complexes of PARP1 with 5′[$^{32}$P]labelled AP-NCP. Lane 1: 147 nt DNA, lane 2: native NCP, lanes 3: AP-NCP, lanes 4–10: complexes of PARP1 with AP-NCP. (I) EMSA analysis of the protein–nucleic acid complexes of PARP2 with 5′[$^{32}$P]labelled native NCP. Lane 1: 147 nt DNA, lane 2: native NCP, lane 3: AP-NCP, lanes 4–14: complexes of PARP2 with AP-NCP.
