## Supplementary material for "The contribution of PARP1, PARP2 and poly(ADP-ribosyl)ation to base excision repair in the nucleosomal context": Figure S3

### SUPPLEMENTARY INFORMATION

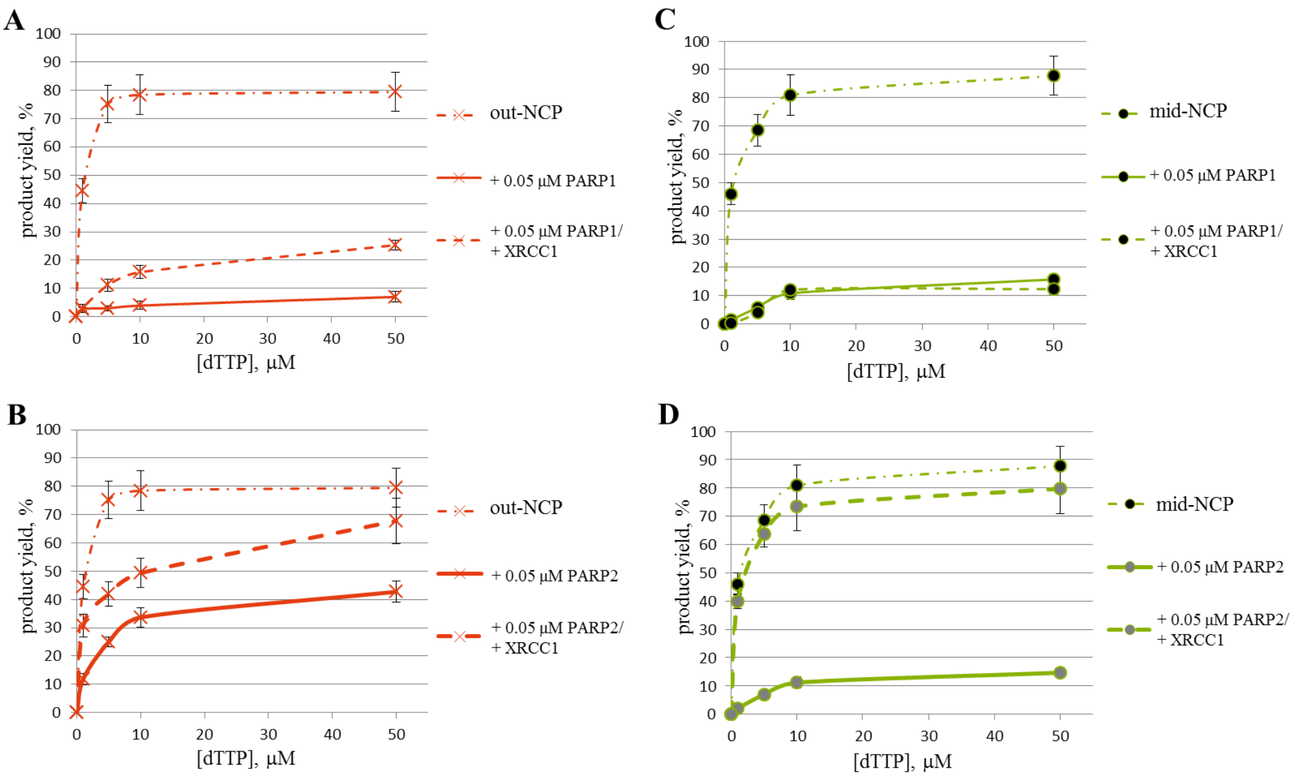


**Figure S3.** The activity of Polβ towards substrates gap-NCP by itself (dot-and-dash red or yellowish-green curves) and in the presence of PARP1, PARP2 and XRCC1. The kinetic curves of dTMP incorporation were obtained using outward-oriented (A, B) or middle oriented (C, D) gap-NCP in the presence of PARP1 (solid red or yellowish-green curves), PARP1 and XRCC1 (dashed red or yellowish-green curves), PARP2 (solid red or yellowish-green curves), PARP2 and XRCC1 (dashed red or yellowish-green curves). For details see the 'Materials and Methods' section.
