## Supplementary material for "The contribution of PARP1, PARP2 and poly(ADP-ribosyl)ation to base excision repair in the nucleosomal context": Figure S4

**SUPPLEMENTARY INFORMATION**

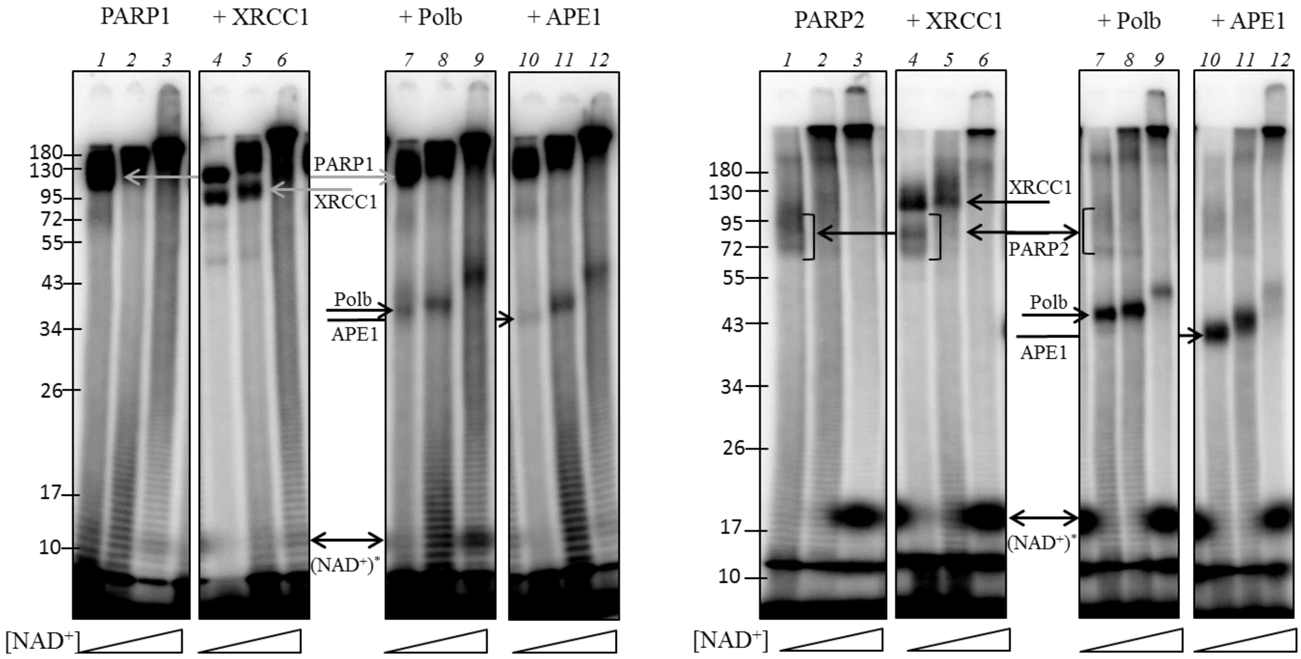


**Figure S4.** PARylation of XRCC1, Polβ and APE1 by PARP1 (left panel) and by PARP2 (right panel) using [$^{32}$P]labelled $NAD^+$. Lanes 1–3: autoPARylation of PARP, lanes 4–6: PARylation of XRCC1, lanes 7–9: PARylation of Polβ, lanes 10–12: PARylation of APE1. The molecular masses in kilodaltons are indicated on the left. The reactions were started by the addition of 1, 10 or 100 μM $NAD^+$ with isotopic dilution of 1:0, 1:9 or 1:99, respectively, towards the solution of 0.1 μM 34 nt 5′-phosphorylated one-window gapped DNA duplex in complex with 0.5 μM PARP1 or PARP2 and 1 μM XRCC1, Polβ or APE1 in reaction buffer with 2 mM $MgCl_2$. The reaction was allowed to proceed for 15 min at 37°C and was stopped by the addition of Laemmle buffer. The products were separated by 12% polyacrylamide gel electrophoresis, dried and subjected to autoradiography using the Typhoon imaging system (GE Healthcare Life Sciences) and analysed in the Quantity One software (Bio-Rad).
