## Supplementary material for "The contribution of PARP1, PARP2 and poly(ADP-ribosyl)ation to base excision repair in the nucleosomal context": Figure S5

### SUPPLEMENTARY INFORMATION

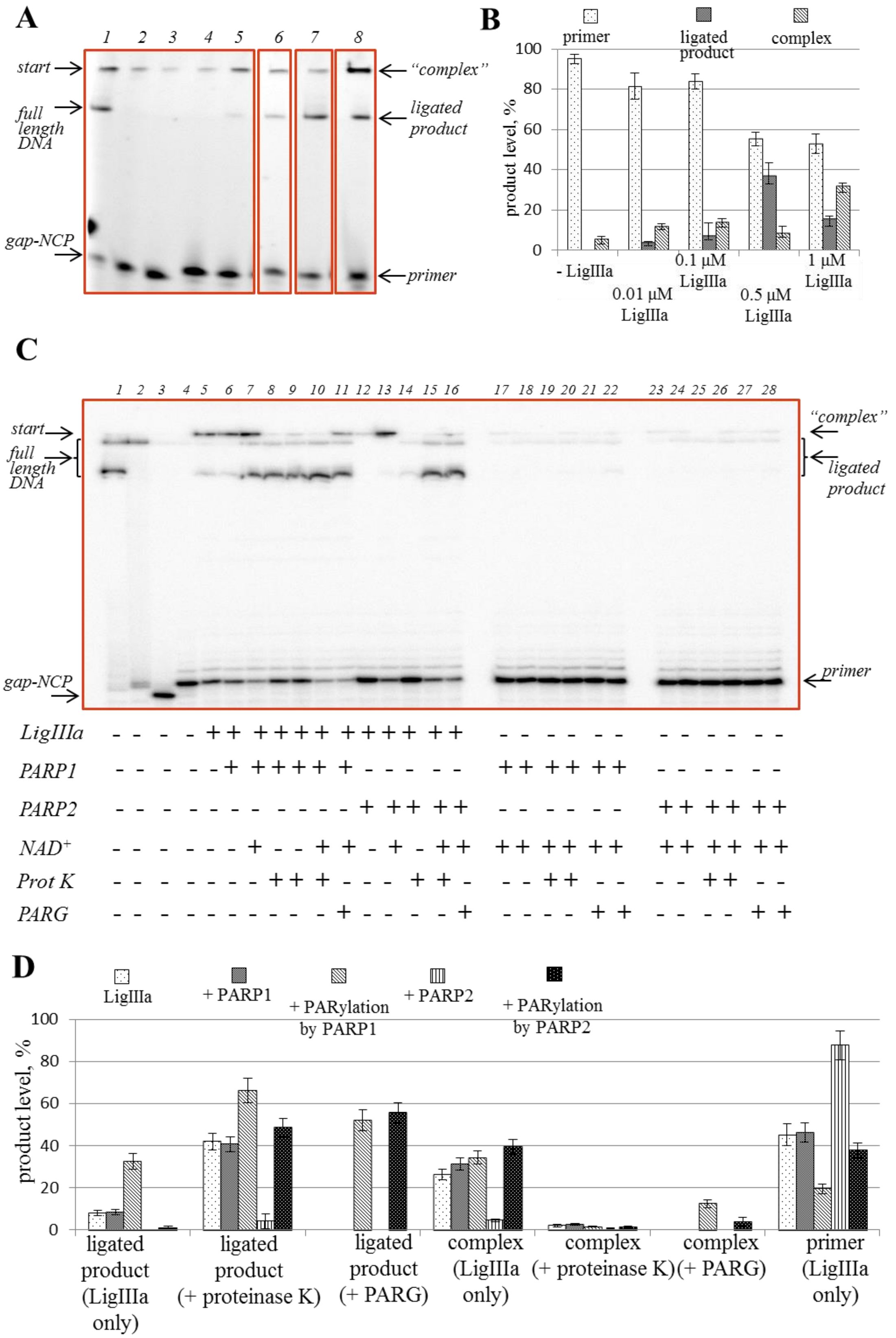


**Figure S5.** The dependence of the LigIIIα activity on substrate 'nicked out-NCP'. Product separation (A) and quantitative analysis (B) of the sealing reaction at different LigIIIα concentrations. The reaction procedures are described in the 'Materials and Methods' section. (A) Lane 1: native out-NCP; lane 2: out-NCP incubated with UDG; lane 3: out-NCP incubated with UDG and APE; lane 4:

out-NCP incubated with UDG, APE, Polβ and dTTP resulting in nicked out-NCP; lanes 5–8: nick sealing in the presence of 0.01, 0.1, 0.5 or 1 μM of LigIIIα. (B) The bars correspond to the level of the initial substrate (dotted), ligated products (hatched) and undenatured complexes (striped) in lines 4–8, respectively.

The product separation (C) and quantitative analysis (D) of the sealing of nicked out-NCP by LigIIIα with PARP1 or PARP2 and PARylation in the presence of XRCC1. The reaction procedures are described in the 'Materials and Methods' section. (C) Lane 1: native out-NCP; lane 2: out-NCP incubated with UDG; lane 3: out-NCP incubated with UDG and APE; lane 4: out-NCP incubated with UDG, APE, Polβ and dTTP resulting in nicked out-NCP; lanes 5–16: nick sealing in the presence of 0.5 μM LigIIIα (lanes 5, 8) and 0.1 μM PARP1 (lanes 6, 9) or PARP2 (lanes 12, 14) without or with 100 μM $NAD^+$ (lanes 7, 10, 11 and 13, 15, 16, respectively). Lanes 8–10, 14–15, 19–20 and 26–26 correspond to lanes 5–7, 12–13, 17–18 and 23–24 with additional treatment with Proteinase K. Lanes 11, 16, 21–22 and 27–28 correspond to lanes 7, 13, 17–18 and 23–24 with additional treatment with PARG. Lanes 17–22: reaction mixtures containing nicked NCP with 0.1 μM PARP1 and 1 or 100 μM $NAD^+$ in pairs. Lanes 23–28: reaction mixtures containing nicked NCP with 0.1 μM PARP2 and 1 or 100 μM $NAD^+$ in pairs. (D) The bars correspond to the level of the indicated product obtained in the reactions with LigIIIα (dotted) only or in the presence of PARP1 (hatched) or PARP2 (vertically stripped) or under PARylation by PARP1 (striped) or by PARP2 (black with white dots). All the data are presented as an average of at least three independent experiments.
