## Supplementary_Table2 for "The contribution of PARP1, PARP2 and poly(ADP-ribosyl)ation to base excision repair in the nucleosomal context"

**SUPPLEMENTARY INFORMATION**

Table S1. *The affinity of PARP1 and PARP2 for native, AP-NCP or gap-NCP according to the EMSA*

$K_d$, nM, for substrates with an outward-oriented lesion

| | NCP | | |
|---|---|---|---|
| | native | AP site | gap |
| PARP1 | 65±3 | 61±5 | 38±4 |
| PARP2 | 192±10 | 130±4 | 57±8 |
